# An adhesion signaling axis involving Dystroglycan, β1-Integrin and Cas adaptor proteins regulates the establishment of the cortical glial scaffold

**DOI:** 10.1101/2022.08.02.502565

**Authors:** Wenny Wong, Jason A. Estep, Alyssa M. Treptow, Niloofar Rajabli, Jennifer N. Jahncke, Teresa Ubina, Kevin M. Wright, Martin M. Riccomagno

## Abstract

The mature mammalian cortex is composed of six architecturally and functionally distinct layers. Two key steps in the assembly of this layered structure are the initial establishment of the glial scaffold and the subsequent migration of post-mitotic neurons to their final position. These processes involve the precise and timely regulation of adhesion and detachment of neural cells from their substrates. Although much is known about the roles of adhesive substrates during neuronal migration and the formation of the glial scaffold, less is understood about how these signals are interpreted and integrated within these neural cells. Here, we provide *in vivo* evidence that Cas proteins, a family of cytoplasmic adaptors, serve a functional and redundant role during cortical lamination. *Cas* triple conditional knock-out (*Cas* TcKO) mice display severe cortical phenotypes that feature cobblestone malformations. Molecular epistasis and genetic experiments suggest that Cas proteins act downstream of transmembrane Dystroglycan and β1-Integrin in a radial glial cell-autonomous manner. Overall, these data establish a new and essential role for Cas adaptor proteins during the formation of cortical circuits and reveal a signaling axis controlling cortical scaffold formation.

## Introduction

The cortex is an exquisitely organized structure and provides an excellent model for understanding the cellular and molecular mechanisms that direct neuronal migration, lamination and stratification [1,2,3,4,5,6,7,8,9]. The mammalian neocortex is composed of six architecturally and functionally distinct layers [10]. Layers II to VI of the cortex develop in an inside first-outside last manner, i.e. Layer VI is born first and Layer II is born last [5,6,11,12,13]. Excitatory neurons that form these layers are born in the ventricular zone (VZ) of the dorsal telencephalon and migrate radially towards the pial surface [2,6,14,15]. After an initial phase of glial-independent migration, the majority of these newly born neurons will switch from a multipolar to a bipolar morphology near the intermediate zone (IZ), and associate with Radial Glial Cell (RGC) processes [10,16 17,18,19,20]. The RGC basal process dynamically changes from highly branched to club-like during early development to establish a scaffold for neuronal migration [10,21,22]. After this initial remodeling event, RGCs have a bipolar shape with an oval nucleus, a short apical process at the VZ, and a long basal process contacting the cortical surface [21]. Neurons use these glial processes as tracks to migrate into the expanding cortical plate (CP) by glia-guided locomotion [3,4,23,24]. Once neurons reach the cortical marginal zone (MZ), they detach from the glial fibers and perform somal translocation to their final location [4,24,25,26]. Thus, two key steps in the assembly of the cortical layers are the establishment of the glial scaffold and the migration of post-mitotic neurons to their final positions [5,21,27].

During these events, neurons and RGCs actively interact with each other and the extracellular matrix (ECM). These processes involve the precise and timely regulation of adhesion and detachment of neural cells from their substrates [28,29,30]. Newborn excitatory neurons constantly remodel their Integrin Adhesion Complexes (IAC) as they migrate radially [29,30,31,32,33,34,35]. In addition, these neurons change adhesive preferences during their migratory path when switching between glial-independent translocation and glial-dependent locomotion [4,19,24]. RGCs remodel their points of contacts and actively engage the ECM as the cortex grows [3,5,21,36,37,38,39,40,41]. Thus, interactions between the basal lamina and the RGC endfeet also play a key role during migration and lamination. In support of this, β1-Integrin and Dystroglycan (Dag1), two important transmembrane regulators of IAC establishment and remodeling, have been shown to be required for cortical histogenesis by acting primarily during the formation and maintenance of the glial scaffold [42,43,44,45,46,47]. Furthermore, mutations that disrupt the expression or glycosylation of Dystroglycan cause cobblestone cortex phenotypes in both human and mouse [46,47,48,49,50].

Although much is known about the roles of adhesive substrates and their cognate receptors during neuronal migration and the formation of the glial scaffold, less is understood about how these signals are interpreted and integrated within these neural cells. The Cas family of cytosolic adaptor proteins are known to participate in IAC/focal adhesion turnover, and accumulating evidence suggests that Cas protein phosphorylation and ensuing recruitment of downstream effectors can be modulated by permissive and adhesive cues *in vitro* [51,52,53]. Cas family members have been shown to mediate a variety of biological processes including cell migration and changes in cell morphology *in vitro* [54,55,56,57]. Three of the four mammalian *Cas* genes are expressed in the developing rodent CNS, including the dorsal telencephalon [58,59,60,61,62]. Using a conditional gene targeting approach in the mouse to inactivate these three *Cas* genes, we previously uncovered a cell-autonomous role for Cas adaptor proteins acting redundantly to mediate Integrin signaling during the organization of the retinal ganglion cell layer (GCL) [61]. Our findings suggest that inactivation of Cas proteins disrupts normal ganglion cell migration and cell positioning [61]. Interestingly, recent genetic evidence from our lab demonstrates that Cas proteins also act cell-autonomously to regulate the adhesive preferences of mammalian Dorsal Root Ganglia (DRG) sensory axons [62], and a similar role has also been shown in epidermal cells [63]. Taken together, these raise the intriguing possibility that Cas proteins could be acting to regulate cortical stratification.

Here, we show that Cas proteins are essential for normal cortical migration and lamination. We also provide evidence suggesting that these adaptor proteins act in a glial-autonomous manner downstream of β1-Integrin and Dystroglycan. These data thus uncover an adhesive signaling pathway essential for cortical glial scaffold formation and lamination.

## Results

As mentioned above, three *Cas* paralogs are expressed during vertebrate development: *p130Cas/BCar1*, *CasL/Nedd9* and *Sin/Efs* [58,59,60,61,62]. A fourth family member, *Cass4,* initially believed to be a pseudogene, appears to only be expressed in the adult [58,64]. We started by undertaking expression analysis of *p130Cas, CasL* and *Sin* during cortical development using multiplex RNAscope *in situ* hybridization [65]. Examination of the RNAscope probes in wild-type (WT) mice at Embryonic day (E) 12.5 (**Fig 1A**) indicated *p130Cas* transcripts are strongly concentrated in the developing preplate (PP), with weaker expression in other areas of the neocortex. *CasL* transcripts are strongest in the cortical hem (midline) and fainter, but distinctly expressed on the dorsolateral neocortex. In comparison to *p130Cas, CasL* transcripts are less prominent in the PP and are found evenly distributed in the subventricular zone (SVZ), and ventricular zone (VZ). Based on transcript expression localization, *p130Cas* puncta are likely found in postmitotic migrating neurons, while both *p130Cas* and *CasL* are expressed in the proliferative pool of cells in the SVZ and VZ. *Sin* is undetectable at this stage as indicated by the absence of signal on coronal sections of the cerebral hemispheres, even at higher magnification. *Cas* gene expression at E14.5 appears somewhat different from their expression at E12.5 (**Fig 1B**). Noticeably, detection of *p130Cas* transcripts at E14.5 is strongest in the IZ instead of the cortical plate (CP). Across the two timepoints, *p130Cas* and *CasL* expression remained consistent in the SVZ and VZ (**Fig 1A and 1B**). The previously undetectable mRNA transcripts for *Sin* are now clearly localized to the SVZ and VZ. The overlap of *p130Cas, CasL,* and *Sin* transcripts in the proliferative zone suggest a possible function for these genes during cortical neurogenesis.

**Fig. 1.**
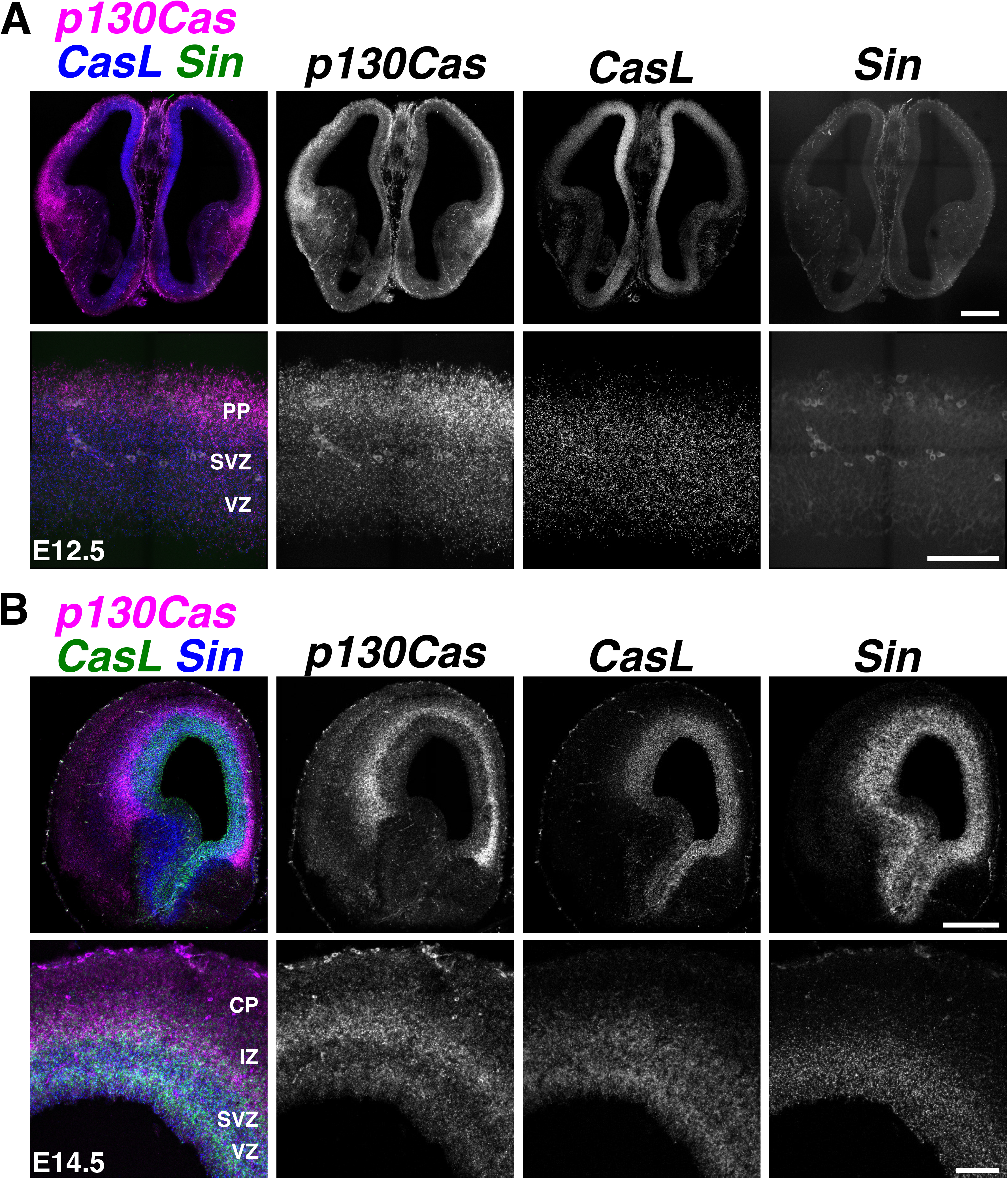
Expression analysis of *Cas* family members. **(A-B)** Expression analysis of *p130Cas* (magenta), *CasL* (**A**, blue; **B**, green), and *Sin* (**A**, green: **B**, blue) mRNA on coronal sections of E12.5 (**A**) or E14.5 (**B**) WT cortex using RNAscope. PP: preplate; CP: cortical plate; SVZ: subventricular zone; VZ: ventricular zone. Scale bars for A and B (top): 500 µm; A and B (bottom): 100 µm.

We then performed a complementary analysis of Cas expression by immunohistochemistry on a GENSAT BAC transgenic mouse line that expresses Enhanced Green Fluorescent Protein (EGFP) under the control of *p130Cas* regulatory sequences [66,67]. This *p130Cas::BacEGFP* transgenic line allows for the detection of cells expressing the *BCar1/p130Cas* gene [61,62]. Coronal sections from E12.5 and E15.5 *p130Cas::BacEGFP* mice were immunostained for EGFP, and co-stained for Tbr1 to help delineate the PP and CP (**Fig 2A-2C**). No signal is detected when using the same EGFP antibodies on WT tissue (**S1 Fig**). At E12.5, EGFP was strongly expressed in the PP, with moderate expression present in the SVZ and VZ (**Fig 2A)**. At E15.5 robust expression is observed in the IZ and CP (**Fig 2B, 2C)**. EGFP is also abundantly expressed in the cortical white matter and blood vessels. Weaker but specific expression of EGFP was found in the VZ and SVZ. EGFP expression in *p130Cas::BacEGFP* is thus mostly consistent with endogenous *p130Cas* mRNA expression in WT animals, as detected by RNAscope. While p130Cas transcript is moderately expressed in the CP at E14.5, the strong expression of EGFP in the CP at E15.5 likely reflects perdurance of EGFP reporter due to accumulation of the fluorescent protein. Taken together, these expression patterns in the CP and proliferative regions indicate that *Cas* genes might be involved in several aspects of cortical development, including cortical neurogenesis, migration, and lamination.

**Figure 2.**
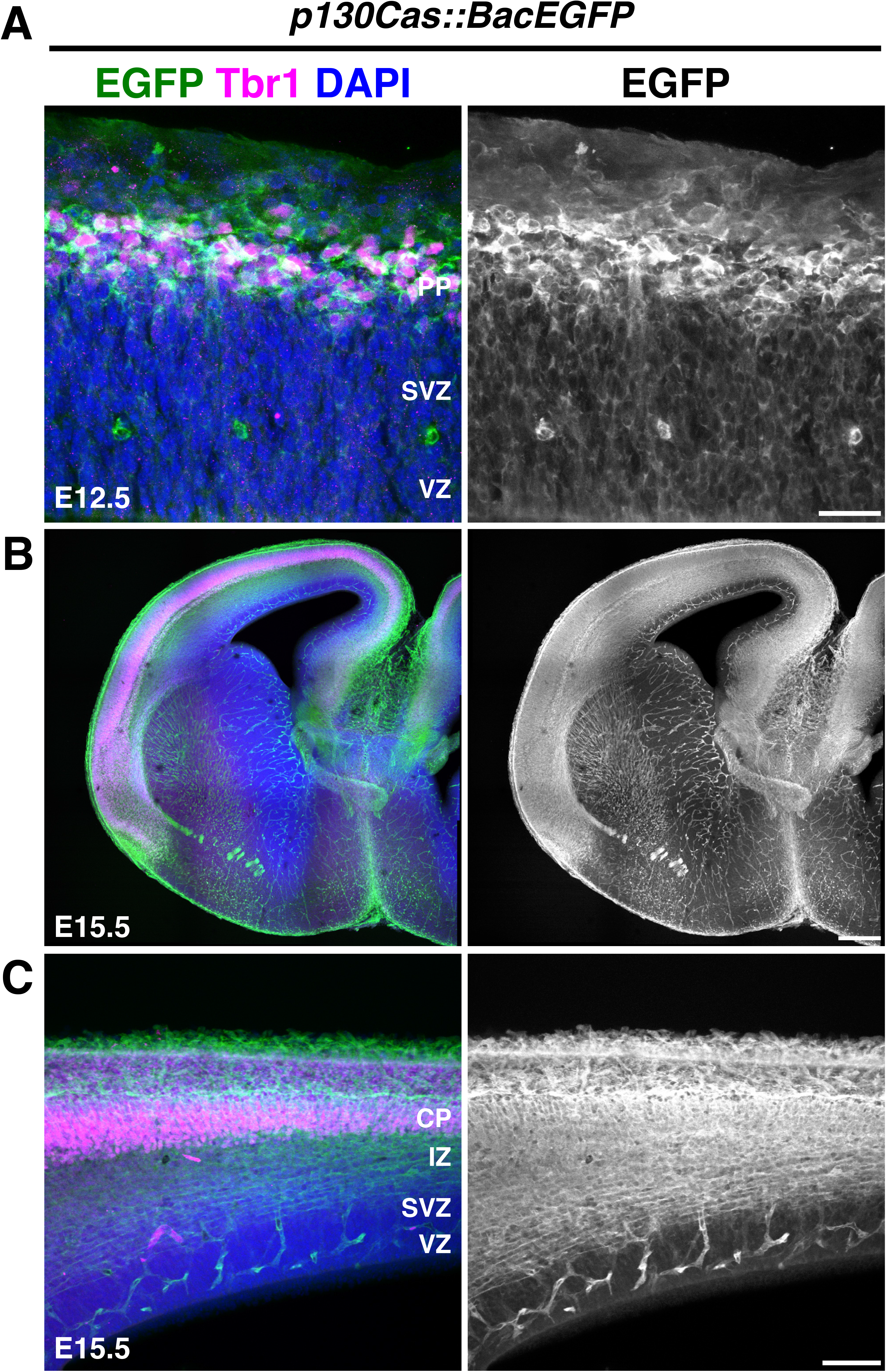
Expression analysis of *p130Cas::BacEGFP* during cortical development. **(A-C)** Coronal sections of E12.5 (**A**) and E15.5 (**B, C**) *p130Cas::BacEGFP* cortices stained for the preplate/cortical plate marker Tbr1 (magenta) and EGFP (green). Nuclei were counterstained with DAPI (blue). PP: preplate; CP: cortical plate; IZ: intermediate zone; SVZ: subventricular zone; VZ: ventricular zone. Scale bars for A: 25 µm; B: 250 µm; C: 75 µm.

To begin to explore whether Cas adaptor proteins play functional roles during cortical circuit assembly, we generated a cortical-specific *p130Cas* ablation in a *CasL^-/-^;Sir^-/-^* double null mutant genetic background (we refer to *p130Cas^flox/-^ ;CasL^-/-^;Sin^-/-^* mice as triple conditional knock-outs: “TcKO”) [61,68,69]. The highly overlapping expression and redundant roles played by Cas proteins make using the *Cas TcKO* mice of vital importance to understand Cas function (**Fig 1A and 1B, 2A-2C**) [61,62]. To drive *Cre* recombinase expression we used the *Em×1Cre* line, which expresses *Cre* in early cortical neural progenitors and radial glial cells (RGCs) in the forebrain [70], resulting in recombination in RGCs themselves and the excitatory neurons in the cortex and hippocampus (**S2A Fig**). RNAscope on E14.5 *Em×1Cre;CasTcKO* coronal sections validated that mutant animals do not produce functional *Cas* transcripts (**S2B Fig**). To test the requirement for *Cas* genes during cortical histogenesis we started by performing an overall examination of *Em×1Cre;CasTcKO* cortical structure using pan-neural markers. When compared to littermate controls (*TcKO*s without *Cre* and *Em×1Cre; p130Cas^+/flox^;CasL^-^’; Sin^-^’* animals), the *Em×1Cre;CasTcKO* adult cortical phenotype appears dramatically different (**Fig 3A**). The smooth cortical surface, typical for a mouse, is replaced with a bumpy or cobblestone surface. Nissl stain, which highlights the contrast between axons and Nissl bodies, shows empty pockets in the cortical plate (**Fig 3A**). A more in-depth analysis of the cortex with a mature neuronal marker (NeuN) indicated the cortex has similar empty pockets as seen in the Nissl stain (**Fig 3B**). Interestingly, NeuN+ cells appear to organize in a wave-like pattern in *Em×1Cre;CasTcKO* animals. In some areas of these cortices, there are cell clusters forming outside of the pial surface and infiltrating into the subarachnoid space of the meninges. This displacement of neurons in the cortex and the cobblestone appearance suggest a possible disruption in the laminar organization of the cortical plate.

**Fig. 3.**
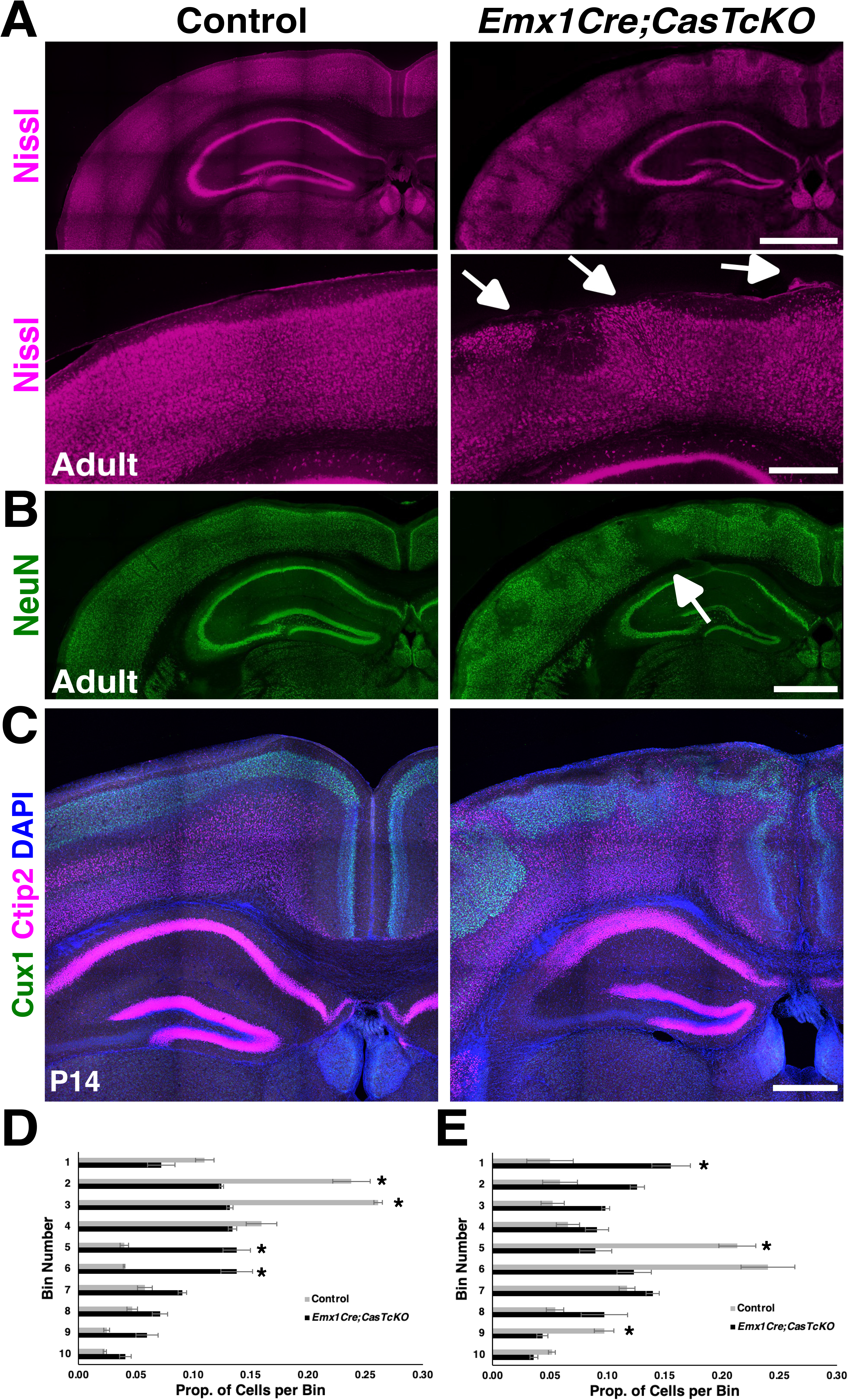
*Cas* genes are required for cortical lamination and neuronal migration. **(A)** Nissl staining of adult control and *Em×1Cre;CasTcKO* coronal cortical sections. Bottom: higher magnification images of Nissl-stained cortices. White arrows mark ectopias. (**B**) Immunostaining of control and *Em×1Cre;CasTcKO* adult coronal sections using the postmitotic neuronal marker NeuN (green) in adult cortices shows a disruption in laminar organization in Emx1Cre;CasTcKO animals. This pan-neuronal marker indicated empty pockets of cells in Emx1Cre;CasTcKO cortices (white arrows). (**C**) Coronal sections of P14 control and *Em×1Cre;CasTcKO* cortices stained for the layer markers Cux1 (green, layer II/III) and Ctip2 (magenta, layer V & VI), counterstained with DAPI (blue). (**D-E**) Quantification of the proportion of Cux1+ (**D**) and Ctip2+ (**E**) cells per bin. Values provided are Mean ± S.E.M., n=3 animals per group, 3 sections per animal, Mann-Whitney U test with Bonferroni correction, *p<0.005 two-tailed test. For data plotted in graphs, see S1 Data. Scale bar for A (top) and B: 1 mm; A (bottom) and C: 500 µm.

The six-layered cortical plate contains distinct populations of projection neurons that can be distinguished based on cell morphology, connectivity, or expression of specific transcription factors [9,13,71]. To test whether the laminar structure of the cortex is affected in *Em×1Cre;CasTcKO* animals, we performed immunohistochemistry with a variety of cortical layer markers. Cux1, Ctip2, Rorβ, and Tbr1 were chosen from a number of transcription factors that maintained their cell lineage identity throughout development [9,13]. Superficial layer neurons (II/III and IV) are identifiable with Cux1 [72], while Rorβ is one of the few markers that will exclusively label layer IV neurons[73]. The deep layer neurons can be identified with Ctip2 (high expression in layer V & low in layer VI) [74] and Tbr1 (layer VI) [75,76]. To precisely quantify any cortical stratification defects, the lamination patterns of control and *Em×1Cre;CasTcKO* animals were analyzed by binning the cortex into ten regions of equal-width as previously described [77,78,79,80,81]. Histological analyses of *Em×1Cre;CasTcKO* animals at P7 and P14 (**Fig 3C-3E; S3A-C Fig**) indicated notable lamination defects for all of the different populations examined in comparison to control animals. Significant differences were observed in the localization of Cux1+, Ctip2+, Rorβ+, and Tbr1+ between P7 *Em×1Cre;CasTcKO* and control animals (**Fig 3D and 3E; S3A-C Fig;** Mann-Whitney *U* test with Bonferroni correction, *p<0.005 two-tailed test for several bins for each marker). The observed cortical dysplasia involves the appearance of Ctip2+/Tbr1+ cell clusters in the upper cortical plate and Cux1+ neurons in the lower cortical plate (**Fig 3C; S3A and S3C Fig**). Furthermore, cortices missing *Cas* genes have a wave-like appearance for layer IV, with Rorβ+ cells invading the superficial cortical plate and meninges (**S3B Fig**). This shows that abnormal positioning of both deep- and superficial-layer neurons contribute to the cobblestone phenotype. This cobblestone phenotype shows a remarkably strong expressivity and 100% penetrance in all postnatal *Em×1Cre;CasTcKO* animals analyzed either by laminar markers or NeuN/Nissl staining, while it was never observed in littermate controls (**Table 1**; n=48 controls, n=46 *Em×1Cre;CasTcKO* animals, Fisher exact test p<0.00001). The number of focal dysplasia per section in P7 *Em×1Cre;CasTcKO* somatosensory cortex averaged 1.47±0.20 dysplasia/mm^2^, while the average was 0 for control animals (**S4 Fig**; Mann-Whitney *U* test with Bonferroni correction, **p<0.0005 n=8 for each genotype). Similar phenotypes are also observed in *Emx1Cre;p130Cas^flox/-^;CasL^-/-^;Sin^+/-^* animals, but not in any other combination of *Cas* family alleles (**S4 Fig,** Mann-Whitney *U* test with Bonferroni correction, *p<0.005 *Em×1Cre;p130Cas^flox/-^;CasL^-/-^;Sin^+/-^* vs control animals, n=3-8 animals per genotype). Despite the misplacement of these laminar markers, the overall density of Ctip2+ and Cux1 + cells were not significantly different between control and *Em×1Cre;CasTcKO* animals (**S3A Fig**; n=5 for each genotype; two tailed t-test *ns*: p=0.12 for Ctip2+ cells; p=0.09 for Cux1+ cells). Taken together these data revealed that *Cas* genes are required for cortical lamination.

The misplacement of layer-specific markers observed in *Em×1Cre;CasTcKO* animals could be due to defects in migration/neural positioning, or defects in neuronal fate specification [9,82,83,84]. To distinguish between these possibilities, newly born cells were tracked during the course of radial migration by performing Ethynyl deoxyuridine (EdU) pulse-chase experiments. This thymidine analog is incorporated in cells undergoing the S-phase of the cell cycle [85] and can be used to observe the final position of cells born at a specific timepoint. EdU was administered to pregnant dams at E12.5 (**Fig 4A**) or E15.5 (**Fig 4B**) to label newly born deep and superficial layer neurons, respectively. Collection at seven days post-intraperitoneal injection for each timepoint (E12.5->P0, E15.5->P3) is sufficient for these populations of excitatory neurons to finish radial migration [7,9,26,39,75,86,87]. Sporadic columns of EdU+ cells are a distinct feature of *Em×1Cre;CasTcKO* animals compared to controls.

**Fig. 4.**
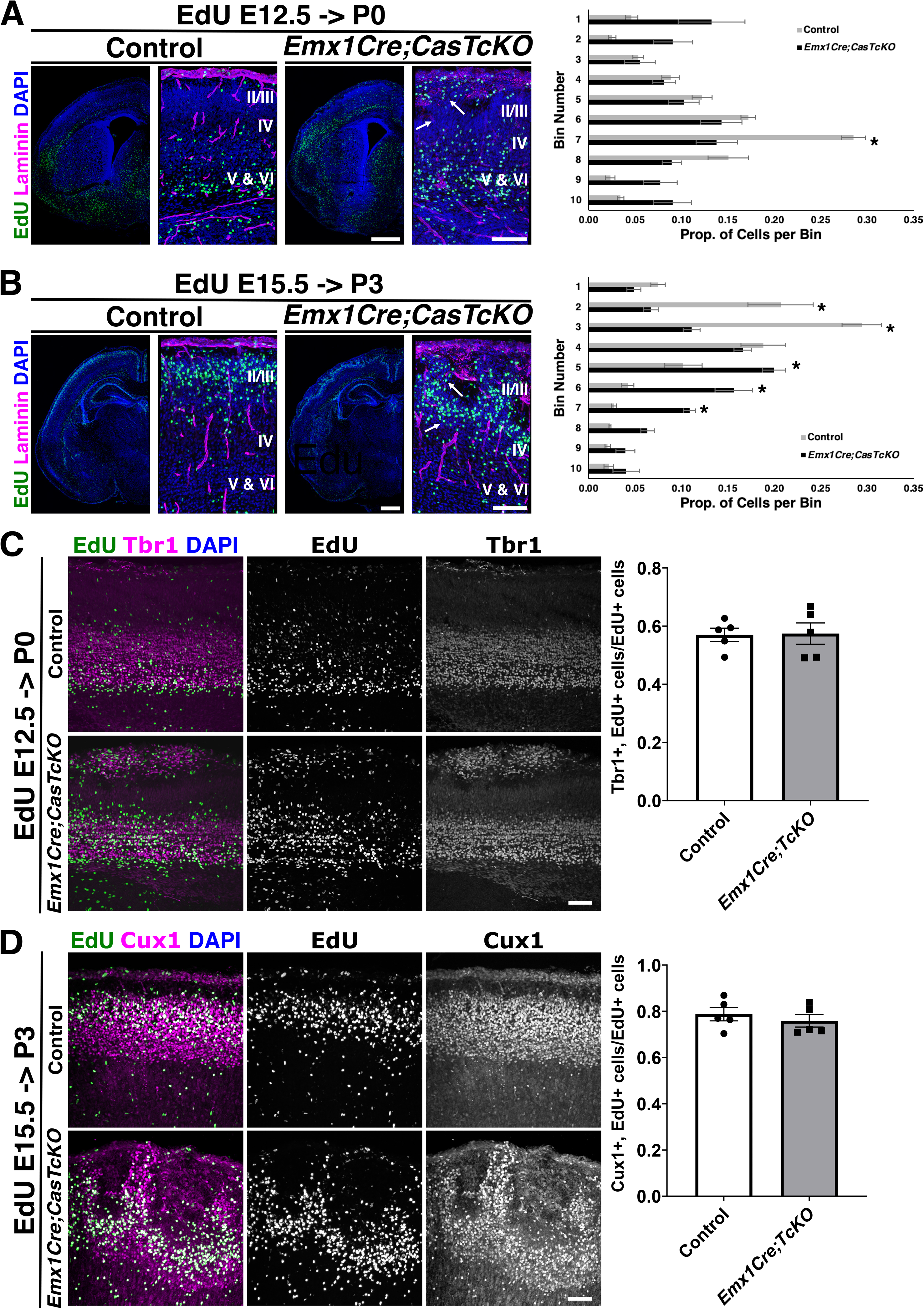
Genetic ablation of *Cas* genes in cortical progenitors results in misplacement of neurons. (**A-D**) Detection of prospective deep layer- (**A, C**) and superficial layer-neurons (**B, D**) labeled by EdU (green) at E12.5 and E15.5, respectively, on coronal sections of control and *Em×1Cre;CasTcKO* cortices. Sections were also stained for laminin (**A, B**), deep layer marker Tbr1 (**C**) or superficial layer marker Cux1 (**D**) shown in magenta, and counterstained with DAPI (blue). (**A, B**) In *Em×1Cre;CasTcKO* animals ectopic deep layer cells are found in the upper CP (**A**, white arrows). EdU also revealed distinct columns of cells extending from the superficial layer into the deep layers (**B,** white arrows) of *Em×1Cre;CasTcKO* animals. Right panels: quantification of proportion of EdU+ cells per bin. n=5 independent samples per group, 2-5 sections per sample, Mann-Whitney U test with Bonferroni correction, *p<0.005 two-tailed test for Bin 7 (**A**) and (**B**) Bins 2, 3, 5, 6 and 7. Bin 1 is the MZ and Bin 10 is ventral to layer VI. (**C, D**) Cells labeled at different timepoints still express the appropriate laminar marker. Right panels: quantification of proportion of EdU+ cells that co-express Tbr1 (**C**) or Cux1 (**D**). n=5 animals per genotype, 3-5 section per animal. Mann-Whitney U Two-tailed test: *ns* p=1 (**C**), *ns* p=0.69 (**D**). Values provided are Mean ± S.E.M. CP: cortical plate; IZ: intermediate zone; SVZ: subventricular zone; VZ: ventricular zone. For data plotted in graphs, see S4 Data. Scale bars for lower magnification panels in A, B: 500 µm; higher magnification panels in A, B: 100 µm. C, D: 75 µm.

Quantification of the position of E12.5 pulsed cells shows a clear trend for EdU+ cells to locate closer to the pial surface (Bin 1) in *Em×1Cre;CasTcKO* cortices than in controls, where they tend to settle in layer VI (**Fig 4A;** Mann-Whitney *U* test with Bonferroni correction, *p<0.005 two-tailed test for Bin 7, n=5 animals per genotype). More noticeable differences are observed when animals are pulsed at E15.5 and collected at P3: prospective superficial neurons that are EdU+ are spread across the ten bins in *Em×1Cre;CasTcKO* animals, whereas in control mice these EdU+ cells are primarily positioned closer to the pial surface (**Fig 4B;** Mann-Whitney *U* test with Bonferroni correction, *p<0.005 two-tailed test for Bins 2, 3, 5, 6 and 7, n=5). This misplacement of cells appears to be caused, at least in part, by columns of ectopic EdU cells in the CP. These data suggest that the cortical dysplasia observed in *Em×1Cre;CasTcKO* animals is likely due to a neuronal mispositioning defect.

To further confirm that the lamination phenotypes are caused by a migration defect and not due to cell fate specification errors, we repeated the pulse-chase experiments but co-labeled the EdU+ neurons with layer specific markers. It is expected that if the cells born at a particular timepoint are misplaced without changing their fate, they should maintain expression of the appropriate cortical layer marker. In control animals 57.0±5% neurons labeled by EdU at E12.5 also co-express the deep layer marker Tbr1 at P0 (**Fig 4C**). A comparable proportion of EdU+ neurons was co-labeled by EdU and Tbr1 in P0 *Em×1Cre;CasTcKO* animals that were treated in the same way (**Fig 4C**; 57.4±8%, Mann-Whitney *U* test with Bonferroni correction, p=1.0 two-tailed test; n=5 for each genotype). When animals were pulsed with EdU at E15.5 and the brains were collected at P3 to label superficial layer neurons, the percentage of EdU+ neurons that co-expressed the superficial layer marker Cux1 was also very similar between control and *Em×1Cre;CasTcKO* mice (**Fig 4D;** 78.7±6.4% vs. 75.9±6% respectively; Mann-Whitney *U* test with Bonferroni correction, p=0.69 two-tailed test; n=5 for each genotype). These results confirm that the EdU+ cells in *Em×1Cre;CasTcKO* mice are properly specified, and thus are most likely mispositioned due to a migration defect.

When is the cortical dysplasia phenotype first observed in the *Em×1Cre;CasTcKO* mutants? By P0 *Em×1Cre;CasTcKO,* animals already display an overt disruption of cortical organization (**Fig 4C**). This disruption of the laminar architecture of the cortex by P0 was confirmed by staining *Em×1Cre;CasTcKO* and control cortices with the cortical marker FoxP2 and the axonal marker L1CAM, suggesting a possible embryonic onset of this phenotype (**Fig 5A**). To visualize dysplasia and ectopias at embryonic stages we labeled E12.5 and 15.5 control and *Em×1Cre;CasTcKO* cortices for Tbr1, which at those stages labels the PP or majority of the CP, respectively. In E12.5 *Em×1Cre;CasTcKO* cortices, localization of Tbr1 + neurons appear indistinguishable from controls (**Fig 5B**). However, by E15.5 ectopic neurons breeching the pial surface are clearly observable in *Em×1Cre;CasTcKO* embryos but are never observed in control animals (**Fig 5C**). By this stage *Em×1Cre;CasTcKO* animals already show ectopic Tbr1+ and Ctip2+ cells positioned outside of the compromised basal lamina (stained with Laminin) (**Fig 5C**-**5E**). On average, at this stage we observed 2.84± 0.3 ectopias/mm of cortical surface length in *Em×1Cre;CasTcKO* embryos, but observed 0 in controls (**Fig 5F**; Mann-Whitney *U* test, *p<0.05 two-tailed test, n=4 animals per genotype). These results establish the developmental onset of the cobblestone phenotype in *Em×1Cre;CasTcKO* cortices between embryonic days 12.5 and 15.5.

**Fig. 5.**
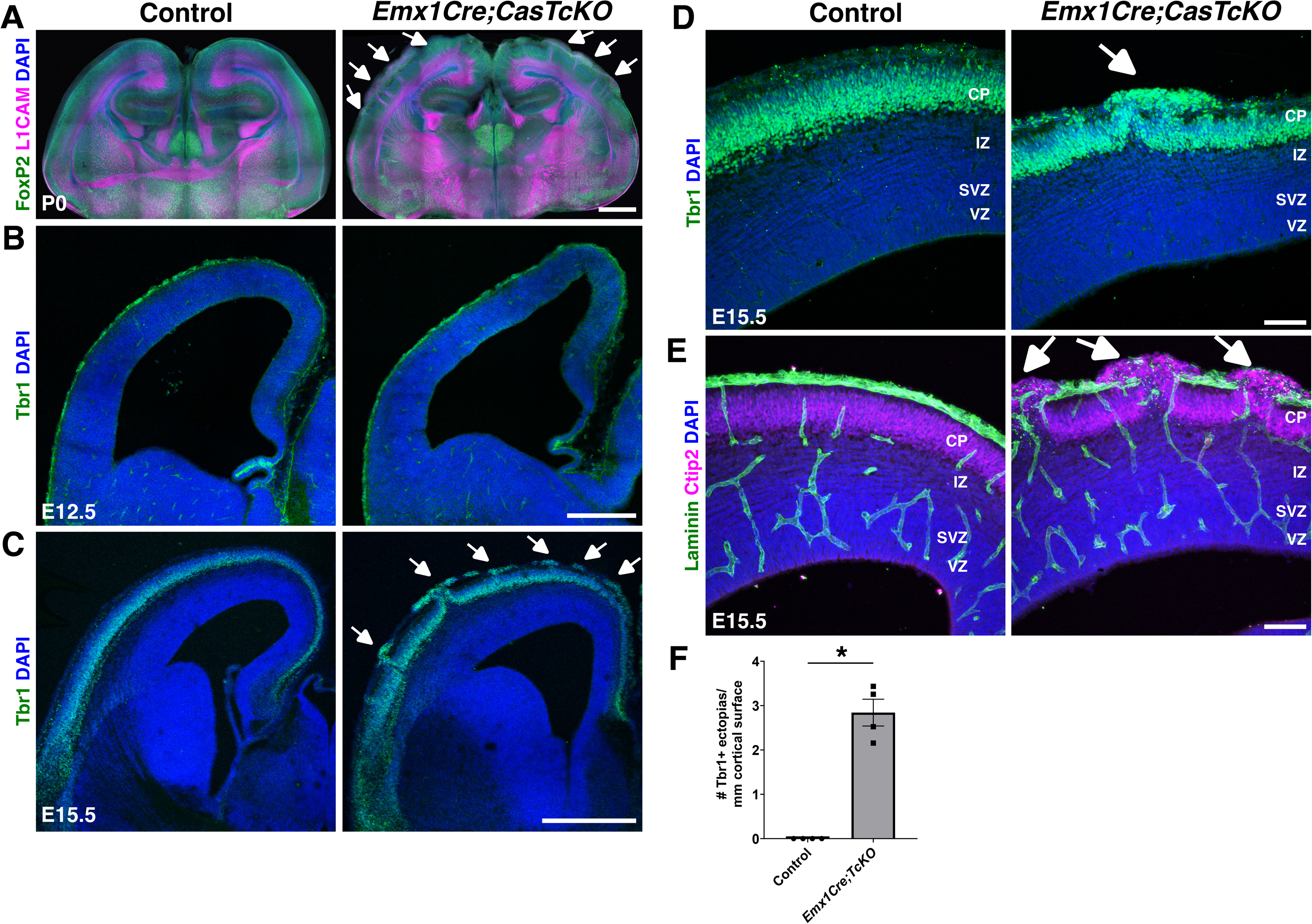
Developmental onset of cortical dysplasia in pancortical *Cas* mutants. **(A-E)** Coronal sections of control and *Em×1Cre;CasTcKO* cortices at P0 (**A**), E12.5 (**B**) and E15.5 (**C-E**) stained for FoxP2 (green) and the axonal marker L1CAM (magenta) (**A**), Tbr1 (green) (**B, C, D**), or Ctip2 (green) and Laminin (magenta) (**E**). While there are no obvious differences observed between *Em×1Cre;CasTcKO* and controls at E12.5 (**B**), by E15.5 there are clear ectopias breeching through the pial surface of *Em×1Cre;CasTcKO* cortices (**C-E**, white arrows). (**F**) Quantification of the number of ectopias per length of cortical surface in mm. Values provided are Mean ± S.E.M. *p<0.05, Mann-Whitney U Two-tailed test, n=4 animals per genotype, 3-5 section per animal. CP: cortical plate; IZ: intermediate zone; SVZ: subventricular zone; VZ: ventricular zone. For data plotted in graphs, see S5 Data. Scale bars: A: 1 mm; B: 250 µm ; C: 500 µm; D and E: 75 µm.

One possible contributing factor for the migration defects observed in animals is the improper splitting of the PP [88,89,90]. This involves the establishment of layer VI within a plexus of pioneer neurons [91,92,93]. Layer VI splits the PP to form the MZ on the surface of the cortex and the subplate (SP) as the ventral boundary. Successive waves of neuronal migration and lamination of cortical neurons form the CP in between these two boundaries. Thus, disruption of this early developmental event can change laminar organization [13,75,94]. To better understand when and how the migration defects observed *Em×1Cre;CasTcKO* animals arise, PP splitting was examined in these mutants and control mice. The use of selected SP and MZ markers at E15.5 indicated a defective PP split in *Em×1Cre;CasTcKO* cortices compared to age-matched littermate controls (**S5 Fig**). Immunostaining for Microtubule Associated Protein 2 (MAP2), which labels differentiating neurons and the cell bodies of SP neurons [75,89], revealed disorganized SP cells underneath the CP. MAP2+ subplate neurons also form ectopic clusters that invade the breached MZ (**S5 Fig**). Staining for Chondroitin sulfate proteoglycan (CSPG), which labels the SP [95], also demonstrated an abnormal distribution of SP neurons, with columns of CSPG+ cells extending dorsally from the SP to contact the pial surface in *Em×1Cre;CasTcKO* cortices. To better characterize PP split phenotypes, we then stained for calcium binding proteins Calretinin and Calbindin (**S5 Fig**). Calretinin is normally expressed by Cajal-Retzius (CR) cells in the MZ and thalamocortical projections in the IZ [96]. Calretinin+ cells in the MZ appear disorganized, forming aggregates on the surface of the cortex, and thalamocortical afferents appear to invade the cortical plate prematurely in *Em×1Cre;CasTcKO* mice compared to controls. This early disorganization of the SP, CP, IZ and MZ was confirmed using Calbindin. Calbindin also labels the MZ and migrating interneurons [97,98]. Calbindin staining shows a disorganized tangential stream of interneurons originating from the medial ganglionic eminence and abnormal positioning of CR cells in the MZ of *Em×1Cre;CasTcKO* cortices. Overall, these suggest that the splitting of the PP and the subsequent organization of the MZ and SP are severely affected in *Em×1Cre;CasTcKO* cortices.

The cobblestone phenotype could be partially caused or compounded by changes in programed cell death during early developmental timepoints. We examined the levels of cleaved (active) Capase-3 in control and *Em×1Cre;CasTcKO* cortices at different developmental stages to visualize the levels of apoptosis before (E12.5), right after (E15.5) and several days after (P0 and P3) the onset of the cortical dysplasia phenotype (**S6A-S6D Fig;** Mann-Whitney *U* two-tailed test, p≥0.7 for all stages, n=3 animals per genotype). The density of caspase-3 activation in control and *Em×1Cre;CasTcKO* mice was not significantly different at any of these stages. These data suggest that programmed cell death is not severely disrupted in *Em×1Cre;CasTcKO* cortices and is unlikely to be a primary cause of the ectopias and cortical dysplasia phenotypes observed in these mice.

The use of the *Em×1Cre* mice precludes us from determining whether *Cas* genes act in a neuronal-autonomous or non-neuronal-autonomous manner during cortical migration and lamination: recombination in these mice occurs as early as E10 in RGCs, the neural progenitors that generate all excitatory neurons and macro glial cells in the dorsal telencephalon [70] (**S2 Fig**). To tease apart the neuronal-autonomous and non­neuronal-autonomous requirements for *Cas* genes, and their relative contributions to the cobblestone phenotypes observed in *Em×1Cre;CasTcKO* animals, we generated *NexCre;CasTcKO* mice*. NexCre* is expressed only in pyramidal cells, but not in RGCs, targeting early postmitotic, premigratory excitatory neurons [99,100]. This pattern of Cre activity was confirmed by crossing the *NexCre* mice to the Cre reporter line *Ai14* [101]. Analysis of *NexCre;Ai14* cortices at E13.5 revealed that tdTomato expression was found in the IZ and CP (**S7A Fig**) and E15.5 included the SVZ [99,100]. Validation of gene inactivation by RNAscope on E16.5 *NexCre;CasTcKO* coronal sections revealed that mutant animals indeed produce next to no *p130Cas* transcripts in the IZ and CP, and indicated a moderate but significant reduction of *p130Cas* transcript levels in the SVZ+VZ (**S7B Fig;** ****P<0.0001 one-way ANOVA w/ Tukey Honestly Significant Difference (HSD) post-hoc test; n=5-6 samples). As expected for the *CasL^--^;Sin^--^* background that these mice are bred into, no mRNA is detected for *CasL* or *Sin* even in controls (**S7B Fig**) [61,68,69].

To elucidate the neuronal-autonomous requirement for *Cas* genes during radial migration and cortical stratification, the laminar organization of *NexCre;CasTcKO* cortices was examined (**Fig 6A; S8A-S8B Fig**). *NexCre;CasTcKO* mutants and control littermates were immunostained with the same layer-specific markers as *Em×1Cre;CasTcKO* animals (**Fig 3C; S3A-C Fig**). A prediction is that, if subpopulations of cortical excitatory neurons are mispositioned in *NexCre;CasTcKO* mice, this will lend evidence to a neuronal-autonomous requirement for *Cas* genes during cortical migration and lamination. There are no apparent differences in lamination between control and *NexCre;CasTcKO* cortices when these are immunostained with a battery of cortical layer markers (**Fig 6A; S8A-S8B Fig**). The transcription factors are expressed in the appropriate laminar layer at P7, where superficial layer (Cux1+), layer IV (Rorβ+), and deep layer (Ctip2+/Tbr1+) subpopulations are spatially distinct. No significant differences in cell positioning were observed for any of the markers examined in control and *NexCre;CasTcKO* animals (p>0.05, Mann-Whitney *U* test with Bonferroni correction). These results strongly suggest that Cas adaptor proteins are not required in a neuronal-autonomous manner during cortical migration.

**Fig. 6.**
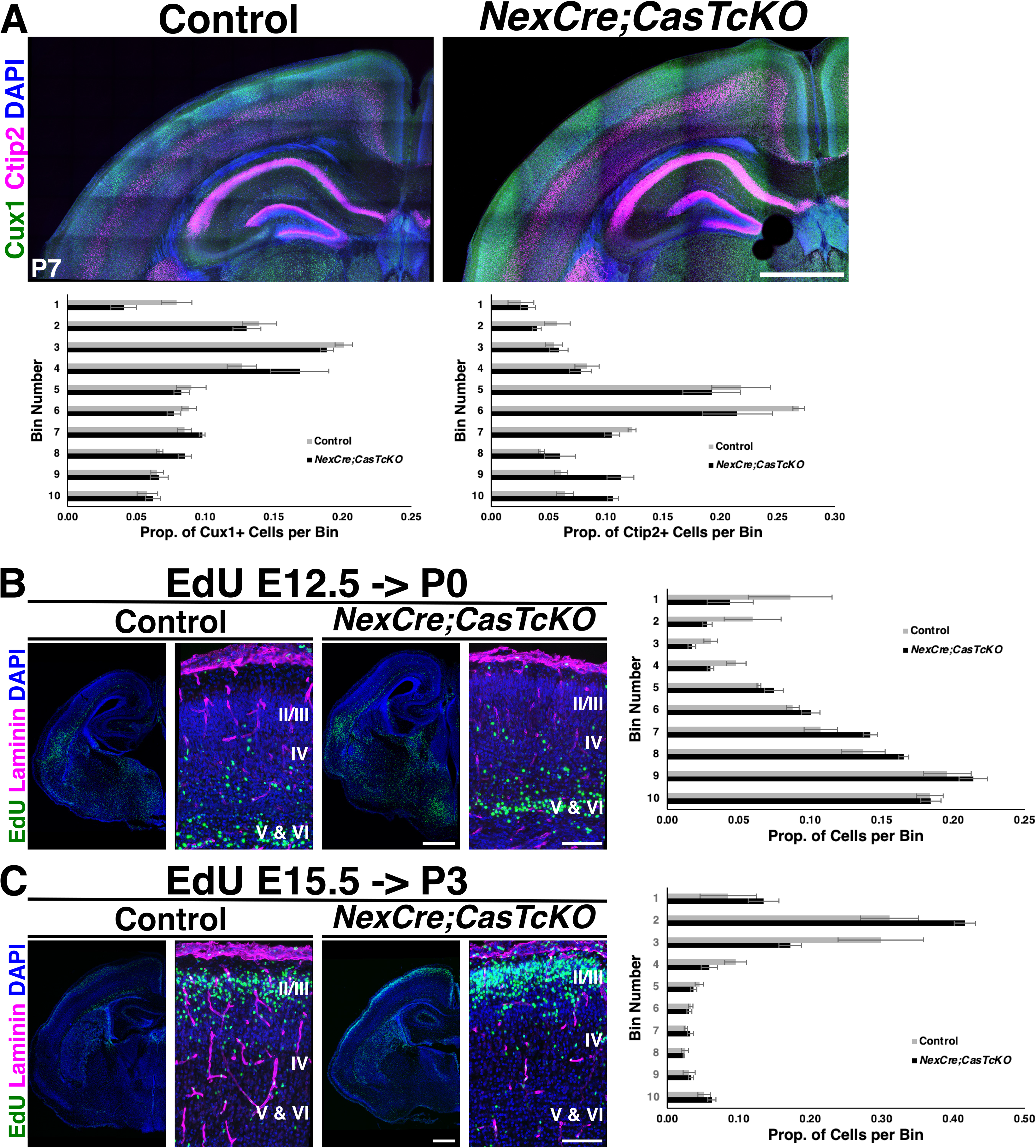
Cas adaptor proteins are required in a non-neuronal-autonomous manner for cortical lamination. (**A**) Coronal cortical sections of P7 control and *NexCre;CasTcKO* mice stained for the layer markers Cux1 (green) and Ctip2 (magenta), counterstained with DAPI (blue). Normal positioning of Cux1+ superficial layer- and Ctip2+ deep layer-neurons. Bottom: quantification of proportion of Cux1+ and Ctip2+ cells per bin. (**B**) Detection of deep layer neurons born at E12.5 and stained for EdU (green; pulsed at E12.5 and collected at P0) in coronal sections of control and *NexCre;CasTcKO* cortices. (**C**) EdU labeling (green) of superficial layer neurons in coronal sections of control and *NexCre;CasTcKO* cortices (pulsed at E15.5 and collected at P3). (**B-C**) Sections were also stained for laminin (magenta) and DAPI (blue). Right panel: quantification of proportion of EdU+ cells per bin. Migration is not affected in *NexCre;CasTcKO* mice. Values given are Mean ± S.E.M., n=3 animals per group, 3 sections per animal, Mann-Whitney *U* test with Bonferroni correction, no significant differences were observed (p>0.05). Bin 1 is the MZ and Bin 10 is ventral to layer VI. MZ: marginal zone; CP: cortical plate; IZ: intermediate zone; VZ: ventricular zone. For data plotted in graphs, see S8 Data. Scale bar for A, lower mag images in B, C: 500 µm; high magnification panels in B, C: 100 µm.

To further confirm the lack of migration defects in *NexCre;CasTcKO,* we performed the same unbiased pulse-chase experiment to track migration of early post­mitotic cells that we used on *Em×1Cre;CasTcKO* animals (**Fig 4A and 4B**). EdU administration into *NexCre;CasTcKO* animals at E12.5 (**Fig 6B**) or at E15.5 (**Fig 6C**) resulted in labeling that was indistinguishable from that of control littermates: both deep and superficial layer neurons exhibit a uniform band of EdU+ cells in the cortical plate (Mann-Whitney *U* test with Bonferroni correction, p>0.05 for all bins examined). Thus, the migration defects observed in *Em×1Cre;CasTcKO* animals are not recapitulated in *NexCre;CasTcKO* mutants. This further demonstrates that *Cas* genes act in a non­neuronal autonomous manner during cortical migration, and that they are likely functioning in RGCs to direct cortical lamination.

As the cortex expands in size, radial glial cells create a scaffold for neurons to migrate [10,9,102]. Interactions between the embryonic pial basement membrane and radial glial endfeet are essential for development of the cortex [21,40]. The absence of the cobblestone appearance and intact basement membrane in *NexCre;CasTcKO* cortices (**Fig 6; S8 Fig**), suggest that the major laminar disruptions observed in the *Em×1Cre;CasTcKO* cortices might be a result of radial glial cell dysfunction. Based on this information, we reexamined the expression of *p130Cas, CasL* and *Sin* in the ventricular zone. To facilitate the labeling of RGC cell bodies and processes we performed *in utero* electroporation (IUE) of a plasmid that drives the expression of EGFP under the Radial Glia-specific *Blbp* promoter [103]. The Blbp-EGFP construct was transfected into WT embryos at E13.5 and tissue was collected at E16.5 to achieve sparse labeling of RGCs. RNAscope analysis of these brains showed that the expression of all *Cas* family members was very similar to that observed at E14.5: *CasL* and *Sin* transcripts were highly enriched in the VZ and SVZ, while *p130Cas* was more broadly expressed from VZ to CP (**Fig 7A** and **7B**). More importantly, RNAscope followed by EGFP immunofluorescence confirmed high levels of expression for *p130Cas* and *Sin* (**Fig 7A**), and *CasL* (**Fig 7B**) mRNA in the cell bodies of RGCs. Little expression, if any, was observed in the RGC endfeet. We next sought to independently validate the expression of *p130Cas* by taking a closer look at EGFP expression in the *p130Cas::BacEGFP* animals. Coronal sections of E12.5 and E15.5 *p130Cas::BacEGFP* brains were immunostained for the RGC marker Nestin, the ECM component Laminin and the intermediate progenitor marker Tbr2 (**Fig 8A-8G**). As mentioned above, EGFP is broadly expressed at E12.5, with moderate expression in the VZ and SVZ that overlaps with Nestin and Tbr2, respectively (**Fig 8A-C)**. At this stage, strongest expression is detected in the PP and basement membrane that are co-labeled by Tbr1 and Laminin, respectively (**Fig 2A** and **Fig 8C**). At E15.5, although EGFP is clearly present in the VZ co-localizing to cells that express Nestin, reporter expression is highest in the IZ, CP and around the basement membrane (Fig **2B** and **8D-G**). Taken together this analysis demonstrates moderate to strong gene expression of *Cas* family members in RGCs, in addition to other cortical cell types.

**Fig. 7.**
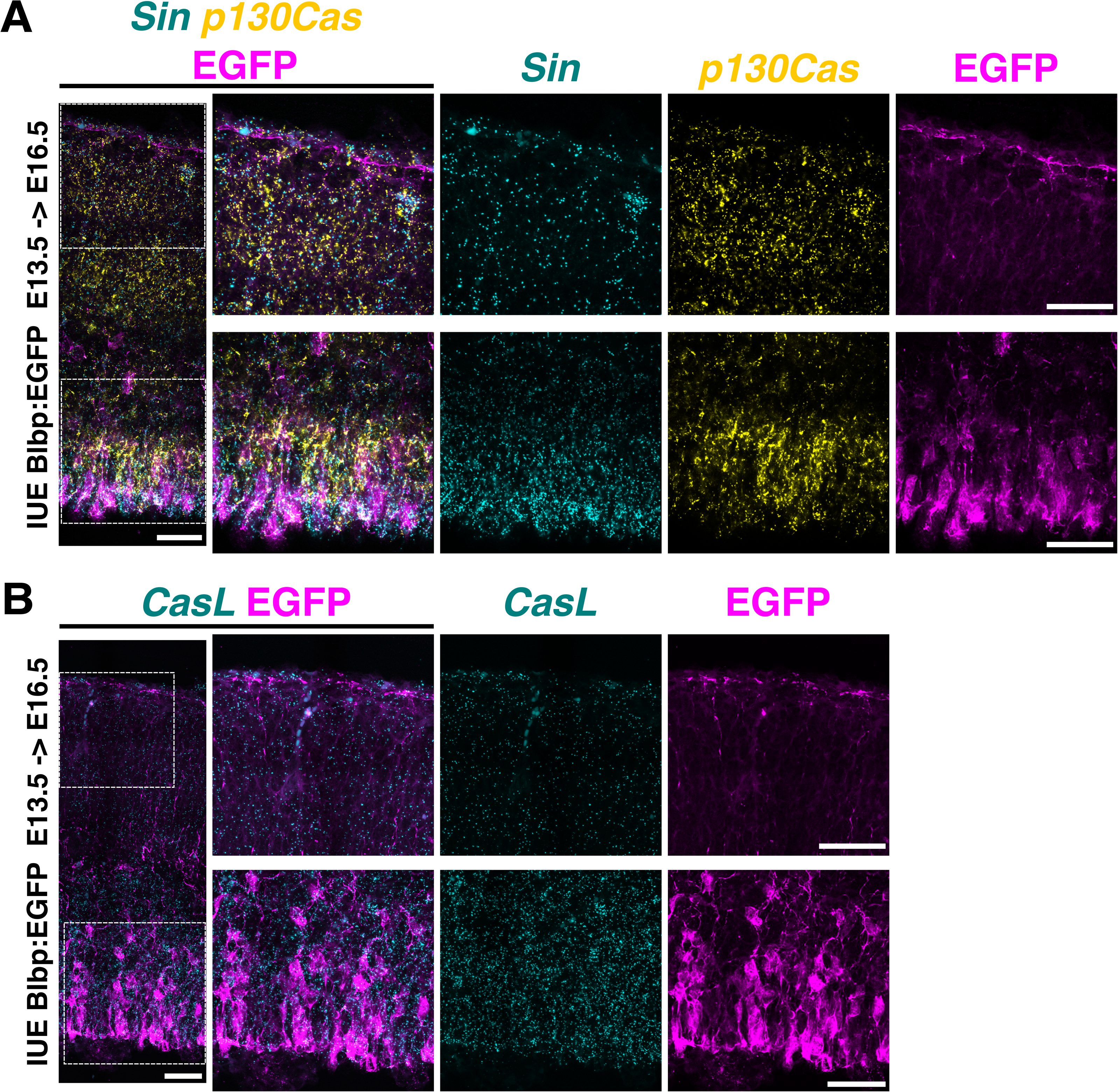
Cas genes are highly expressed in RGCs. **(A-B)** RNAscope for *p130Cas* (yellow) and *Sin* (cyan) mRNA (**A**), or *CasL* (cyan) (**B**) on coronal sections of E16.5 WT embryos electroporated with Blbp:EGFP at E13.5. Sections were also immunostained for EGFP (magenta). Dotted squares in the left panels highlight the regions displayed at higher magnification in the middle and right panels. All *Cas* genes are highly expressed in RGC cell bodies (lower panels), but are less abundant in endfeet (upper panels). Scale bars: 50 µm.

**Figure 8.**
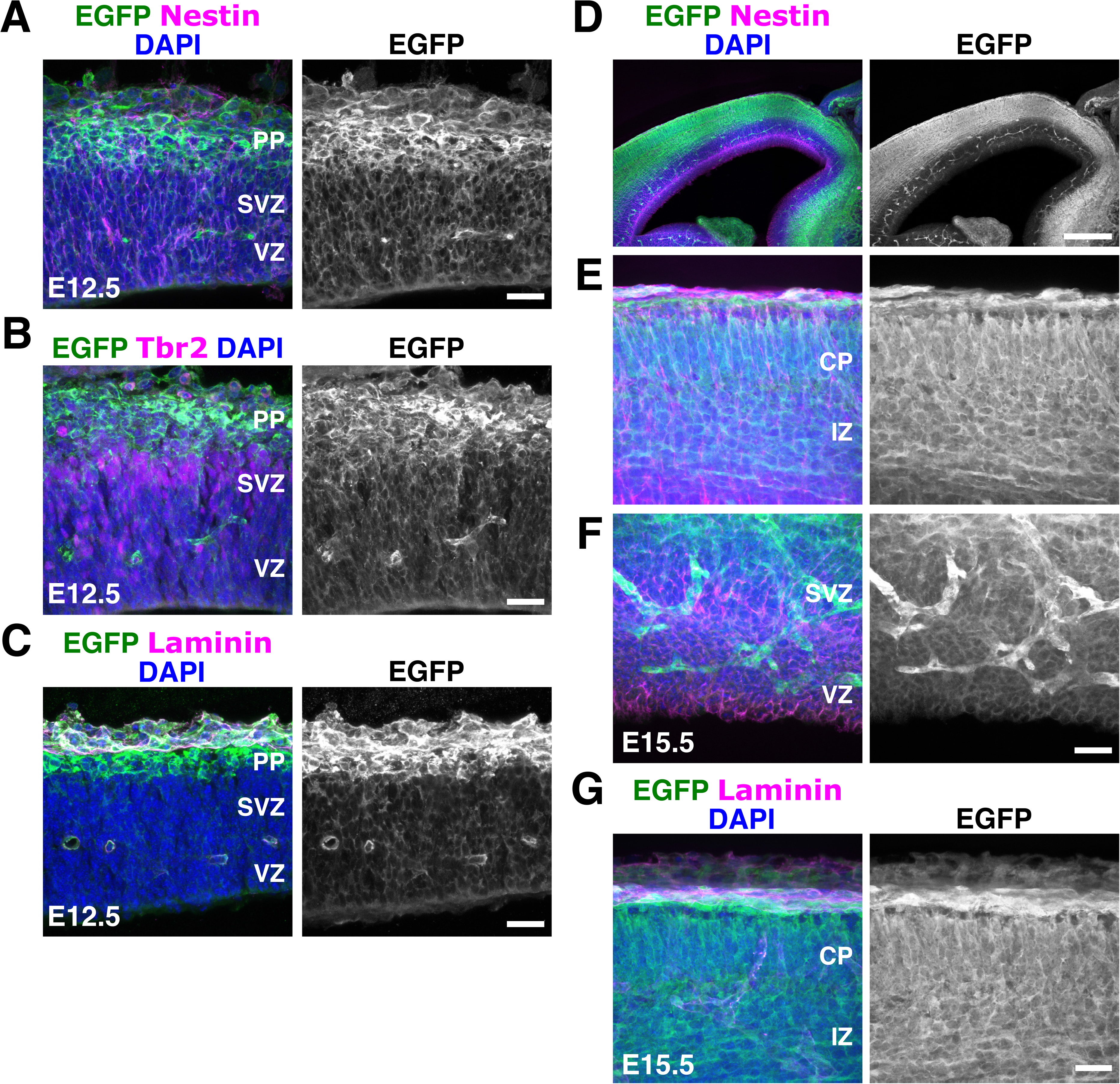
Expression of *p130Cas::BacEGFP* in the VZ and CP. **(A-G)** Coronal sections of E12.5 (**A-C**) and E15.5 (**D-G**) *p130Cas::BacEGFP* cortices stained for the RGC marker Nestin (**A, D, E, F**), Tbr2 (**B**) or laminin (**C, G**) (magenta), and EGFP (green). Moderate expression is observed in the VZ/SVZ, but expression is highest in CP. DAPI was used to counterstain nuclei (blue). PP: preplate; CP: cortical plate; IZ: intermediate zone; SVZ: subventricular zone; VZ: ventricular zone. Scale bars: A, B, C, E,F,G: 25 µm ; D: 250 µm

To test whether *Cas* genes are required for the embryonic pial basement membrane and radial glial endfeet integrity, we performed histological assessment of the cortical scaffold of *Em×1Cre;CasTcKO* at E15.5 (**Fig 9A**). While no obvious defects were observed in control animals, there are several disruptions to the glial-pial interface in *Em×1Cre;CasTcKO* cortices. There is widespread rupture of the basal lamina as indicated by regions with breached Laminin staining. The Nestin+ RGCs do not make proper adhesion contact with the basal lamina at the exposed Laminin sites, where basal processes extend into the subarachnoid space. These results suggest that *Cas* function is required for the maintenance of the embryonic pial basement membrane-RGC interactions.

**Fig. 9.**
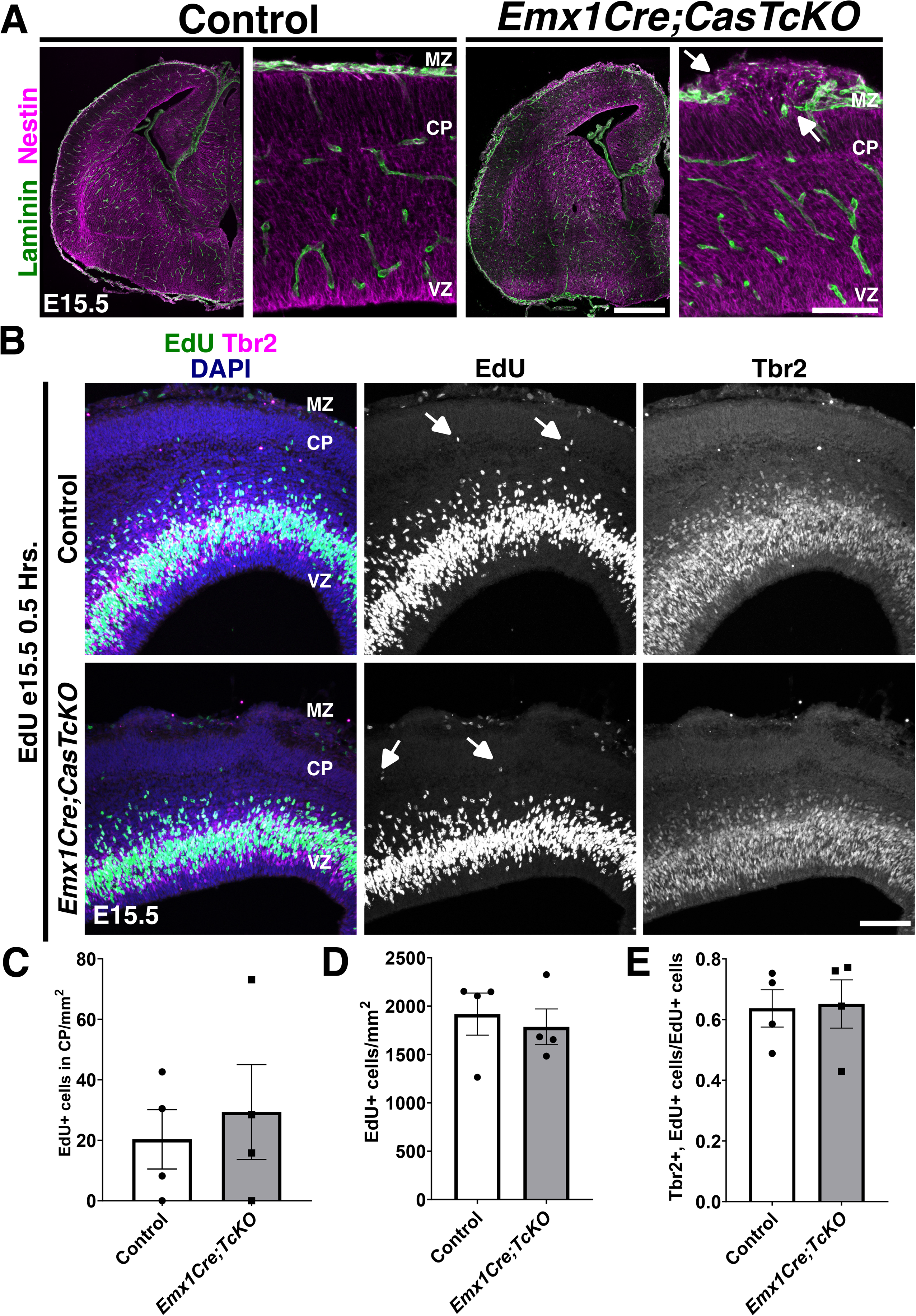
A requirement for Cas adaptor proteins during glial scaffold formation. **(A)** Laminin (green) and Nestin (magenta) antibody staining on E15.5 control and *Em×1Cre;CasTcKO* coronal sections. Radial glial endfeet do not make proper contact with the basal lamina in *Em×1Cre;CasTcKO* animals at exposed laminin sites (arrows). (**B**) Detection of proliferative cells using EdU (green) at E15.5, co-labeled with Tbr2 (magenta). Proliferating cells are occasionally observed within the CP of control and *Em×1Cre;CasTcKO* animals (white arrows). (**C**) Quantification of density of EdU+ cells in CP. Mann-Whitney *U* two-tailed test, p=1, n=4, 4 sections per animal. (**D**) Density of EdU+ cells per area of cortex. Mann-Whitney *U* two-tailed test, p=0.89, n=4 animals per genotype, 3 sections per animal. (**E**) Proportion of EdU+ cells that co-express Tbr2. Mann-Whitney *U* two-tailed test, p=0.49, n=4 animals per genotype, 3-5 sections per animal. Values given are Mean ± S.E.M. CP: cortical plate; MZ: marginal zone; VZ: ventricular zone. For data plotted in graphs, see S10 Data. Scale bars: 100 µm.

To determine whether the disrupted basement membrane-RGC interaction caused mispositioning of proliferating progenitor cells as observed in other models of cortical heteropia and dysplasia [47,104], EdU was again administered to *Em×1Cre;CasTcKO* animals (**Fig 9B; S9 Fig**). However, collection was 0.5 Hrs. following injection at E12.5 (**S9 Fig**) or E15.5 (**Fig 9B**) for these experiments. This short EdU pulse labels the proliferating cells during neurogenesis of deep layer and superficial layer neurons, respectively. We then stained these brains for EdU and the intermediate progenitor marker Tbr2. At E12.5, EdU+ proliferating neural progenitors are positioned in the SVZ and VZ (**S9A Fig**). At this stage, a few proliferative cells are also observed in the PP of both controls and *Em×1Cre;CasTcKO* animals (**S9A and S9B Fig)**. At E15.5, proliferative cells are mainly restricted to the SVZ and VZ of control animals, with a few cells labeled by EdU in the CP and MZ. In *Em×1Cre;CasTcKO* animals, EdU+ cells are also primarily observed in the SVZ and VZ. While occasionally a few EdU+ cells were localized to the MZ and upper CP regions, no significant differences were observed between the density of proliferative cells in the CP of *Em×1Cre;CasTcKO* and control animals (**Fig 9C**). This indicates that the breakage in the basement membrane and disruption of radial glial endfeet observed in *Em×1Cre;CasTcKO* cortices does not result in ectopic positioning of progenitor cells like in *Eml1* or *Dystroglycan* mutants [47,104]. To test whether the overall levels of proliferation and neurogenesis were affected, we measured the density of EdU+ cells in E12.5 and E15.5 control and *Em×1Cre;CasTcKO* cortices (**Fig 9D** and **S9C Fig**). Importantly, the density of proliferating cells is not significantly different between these mutants and controls (Mann-Whitney *U* two-tailed test, p=0.34 for E12.5 and p=0.89 for E15.5, n=4 animals per genotype). To further examine possible defects in neurogenesis and to test whether the transition from RGC to intermediate progenitor is disrupted in *Emx1Cre;CasTcKO* cortices, we quantified the proportion of EdU+ cells that co­expresses Tbr2. This proportion is not significantly different between control and *Emx1Cre;CasTcKO* animals at E12.5 (**S9B and S9D Fig**) or E15.5 (**Fig 9B and 9E;** Mann-Whitney *U* two-tailed test, p=0.49 for both stages, n=4 animals per genotype), suggesting that the disruptions in cortical lamination observed in these mutants are also unlikely to be caused by an early transition from RGC to intermediate progenitor fate.

Dystroglycan (Dag1) provides a structural link between the basement membrane and components that remodel the actin cytoskeleton [46,105]. Glycosylation of the extracellular alpha subunit of Dag1 is responsible for maintaining the pial basement membrane integrity [46,47]. Previous analysis of *NestinCre;Dagl^flox/flox^* animals at E14.5 to P0.5 showed a heterotopic distribution of cortical neurons [47], and an intriguing resemblance to the *Em×1Cre;CasTcKO* cortical phenotype. We thus re-examined these cortical phenotypes at P7 using the same layer markers and the same cortical Cre driver *(Em×1Cre)* that we used to study the *CasTcKO* mice. Genetic ablation of *Dag1* using *Em×1Cre (Emx1Cre;Dag1^flox-^)* indeed caused cobblestone malformations, as suggested by previous studies using a pan-neural Cre [46,47] (**Fig 10A and 10B**). Mispositioning of the different neuronal subpopulations in *Emx1Cre;Dag1^flox-^* mice is strikingly similar to that of *Em×1Cre;CasTcKO* animals, where superficial layer neurons under-migrate and deep layer neurons over-migrate (**Fig 10A and 10B**). As in *Em×1Cre;CasTcKO* animals, the cobblestone cortex phenotype displays strong expressivity and is 100% penetrant (**Table 1**; n=18 for *Emx1Cre;Dag1^flox-^* and controls; Fisher exact test p<0.0001). Interestingly, these phenotypes are also likely RGC-autonomous, as deletion of *Dag1* in postmitotic neurons using *NexCre* does not disrupt cortical lamination [46,106]. In addition to supporting the reproducibility of previous reports, this provides definitive evidence that the *Em×1Cre;CasTcKO* cortical organization phenocopies multiple aspects of the defects observed when *Dag1* is conditionally ablated in early cortical progenitors.

**Fig. 10.**
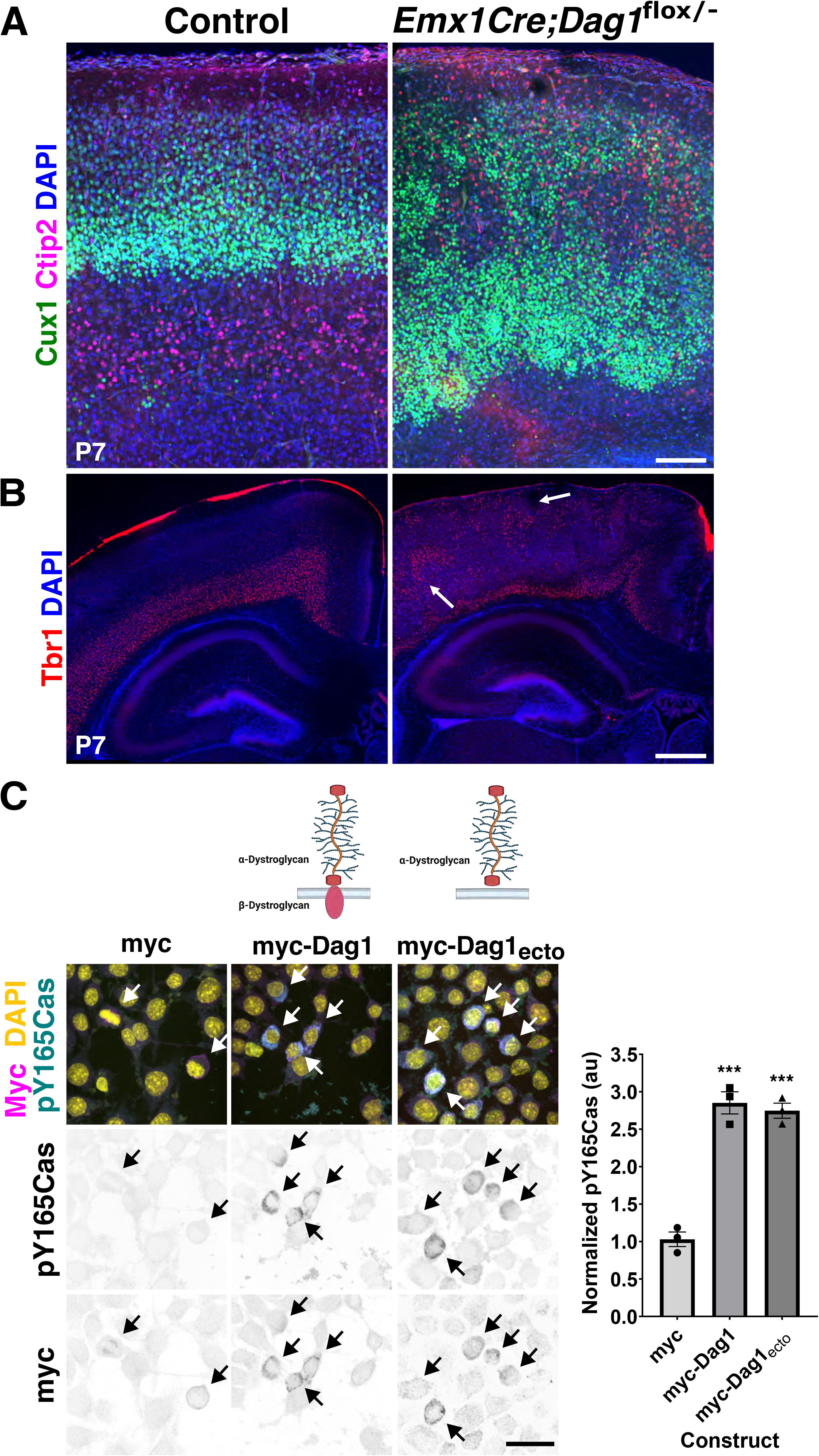
*Dag1* is required for cortical lamination and sufficient to increase Cas phosphorylation in neural cells. (**A-B**) Coronal cortical sections of P7 control and *Emx1Cre;Dag1^flox/-^* animals stained for the layer markers Cux1 (green) and Ctip2 (magenta) (**A**) or Tbr1 (red) (**B**), counterstained with DAPI (blue). *Emx1Cre;Dag1^flox/-^* mice display a similar cortical phenotype to *Em×1Cre;CasTcKO* mutants with disorganized distribution of superficial and deep layer neurons. Scale bars for A: 70 µm; B: 500 µm. (**C**) The extracellular domain of Dag1 is sufficient to increase Cas phosphorylation. Immunofluorescence for Myc (magenta) and pY165Cas (cyan) of Neuro2A cells transfected with control vector (myc), full-length Dag1 (myc-Dag1) or the extracellular domain of Dag1 (myc-Dag1_ecto_). DAPI was used to stain nuclei (yellow). Both myc-Dag1 and myc-Dag1_ecto_ cause a significant increase in pY165Cas. Values provided are Mean ± S.E.M, n=11-23 cells from 3 independent experiments; ANOVA, p<0.0001. Tukey HSD *** p<0.0001 myc-Dag1 vs myc, and myc-Dag1_ecto_ vs. myc. No significant difference was observed between myc-Dag1 and myc-Dag1_ecto_. For data plotted in graphs, see S12 Data. Scale bars for A: 70 µm; B: 500 µm; C: 25 µm.

As mentioned above, Dag1 is an adhesion receptor that provides a direct link between the ECM and pathways involved in cytoskeletal remodeling [107]. While the extracellular domain of α-Dag1 interacts with ECM components, the cytoplasmic domain of β-Dag1 can function to regulate ERK/MAPK and Cdc42 pathway activation, and even directly interact with actin binding proteins [108,109,110,111]. Given the phenotypic similarities observed in *Em×1Cre;CasTcKO and Emx1Cre;Dag1^flox/-^,* we asked whether Dag1 could be modulating Cas activity by promoting/enhancing its phosphorylation to effect downstream signaling. We started by overexpressing myc-tagged full-length Dystroglycan (myc-Dag1) in the neural cell line Neuro2A and compared the levels of tyrosine phosphorylated p130Cas (pY165Cas) to that of cells transfected with a myc-tagged empty vector alone. Transfection of myc-Dag1 increased pY165Cas levels by over 2.5 fold compared to cells transfected with control vector (**Fig 10C**; 1.03±0.16 au vs. 2.85±0.25 au; One-way ANOVA p<0.001, Tukey HSD test ***p<0.0001). Based on the ability of Dag1 to serve as a scaffold for ERK signaling components, we hypothesized that the intracellular domain of Dag1 could be required for Dag1-induced Cas phosphorylation. To test this hypothesis we overexpressed the extracellular domain of Dag1 (myc-Dag1_ecto_) and compared p130Cas phosphorylation levels vs control- and full-length Dag1-transfected cells. Surprisingly, myc-Dag1_ecto_ increased pY165Cas levels notably and significantly compared to control (**Fig 10C**; 2.74±0.17 au; One-way ANOVA p<0.001, Tukey HSD test ***p<0.0001). The increase in pY165Cas was not significantly different than the one observed after full-length myc-Dag1 transfection (Tukey HSD test ns, p>0.05). This suggests that both full-length and a truncated form of Dag1 lacking a cytoplasmic domain are sufficient to induce an increase in p130Cas phosphorylation in a neural cell line.

The intracellular domain of Dag1 appears to be dispensable for the Dag1-dependent increase in Cas phosphorylation in vitro. Is the cytoplasmic domain of Dag1 required for cortical lamination? To answer this question we examined the cortical lamination pattern of a knock-in mouse line in which the endogenous *Dag1* coding sequence was replaced with a truncated form of Dag1 that lacks the intracellular domain *(Dag1^βcyto^),* rendering it unable to bind dystrophin/utrophin or initiate ERK/MAPK or Cdc42 signaling [112]. We performed this analysis using similar markers to those employed for characterizing the *Em×1Cre;CasTcKO* and *Emx1Cre;Dag1^flox/-^* cortical phenotypes. Surprisingly, in *Dag1^βcyto/-^* all the layer markers are expressed in the appropriate pattern in all postnatal animals examined, where superficial layers (Cux1+), layer IV (Rorβ+), and deep layer neurons (Ctip2+ and Tbr1+) are clearly delineated (**Fig 11A-C**). Furthermore, no evidence of ectopias or cortical dysplasia was observed in these mutants compared to littermate controls (**Fig 11A-C**; **Table 1**; n=10 postnatal animals for both genotypes; Fisher Exact test p=1). These data suggest that the intracellular domain of Dag1 is dispensable for cortical lamination.

**Fig. 11.**
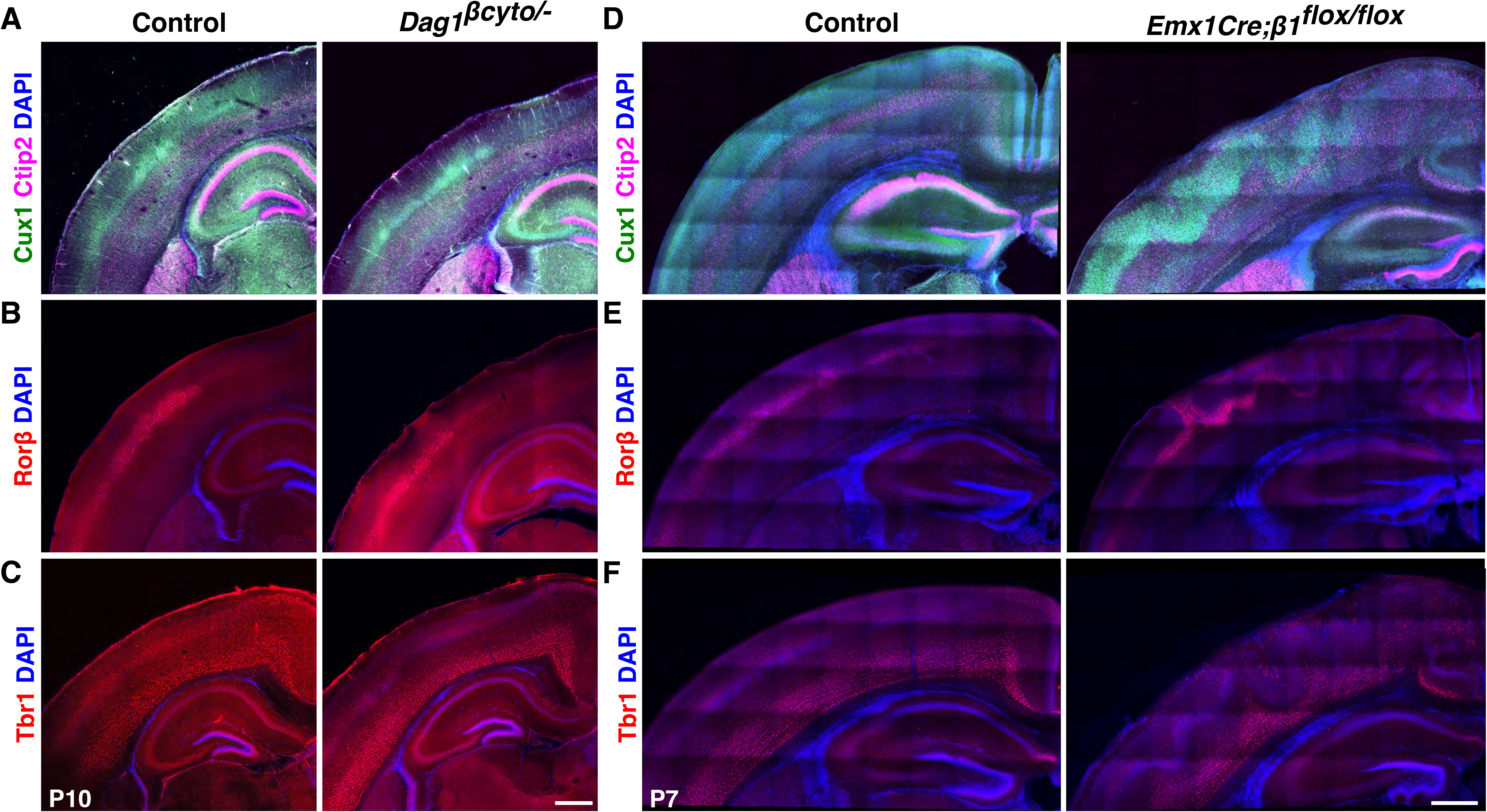
The cytoplasmic domain of Dag1 is not required for cortical lamination, but *β1-Integrin* is essential for this process. (A-C) The cytoplasmic domain of Dag1 is dispensable for cortical lamination. Coronal sections of P7 control and *Dag1^βcyto/-^* cortices immunostained for Cux1 (green) and Ctip2 (magenta) (**A**), Rorβ (red) (**B**) or Tbr1 (red) (**C**). DAPI was used as nuclear counterstain (blue). No obvert lamination defects were observed in these animals (n=10). **(D-F)** *β1-Integrin* ablation results in Cobblestone cortex phenotypes. Immunostaining on coronal sections of P7 control and *Emx1Cre;β1-Integrin^flox/flox^* cortices for Ctip2 (magenta) and Cux1 (green) (**D**), RORβ (red) (**E**) or Tbr1 (red) (**F**). Similar laminar disorganization and cobblestone malformation were observed in *Em×1Cre;CasTcKO* animals. Scale bars: 500 µm.

Several studies using cell-based assays have shown that Dag1 is required for the initial clustering of Laminin-1 on cells [113,114,115,116]. Integrins subsequently bind to clustered Laminin through a distinct interaction site to transduce outside-in signaling [116,117,118]. A previous study showed that Cas phosphorylation in the retina is highly dependent on β1-Integrin function [61]. Interestingly, pan-neural *β1-Integrin* ablation [43] leads to heterotopias similar to the ones observed in *Em×1Cre;Dag1^flox/-^* mice and *Em×1Cre;CasTcKO.* These phenotypes also appear to be RGC-autonomous, since *Ne×Cre-driven* ablation of *β1-Integrin* does not cause these types of defects [45]. The laminar organization of *NestinCre;β1-Integrin* mice has not been fully characterized [43,45]. To begin to probe whether *β1-Integrin* might be acting together with *Cas* genes and *Dag1* during cortical scaffold formation, we revisited the cortical phenotypes of *β1-Integrin* knock-outs using layer-specific markers and *Em×1Cre* as a driver. Staining for layer-specific transcription factors confirmed a distinct cobblestone phenotype in *Emx1Cre;β1-Integrin^flox/flox^* cortices, as compared to control littermates *(Em×1Cre;βl-Integrin^flox/+^)* (**Fig 11D-13F**). Cux1, Ctip2, Rorβ, and Tbr1 labeling revealed that cortical layer misplacement occurred for all the examined subpopulations in *Em×1Cre;βl-Integrin^flox/flox^* animals (**Fig 11D-11F**). The cobblestone phenotype shows a high level of expressivity and 100% penetrance in *Em×1Creβ1-Integrin^flox/flox^* animals but is never observed in control littermates (**Table 1**; n=17 postnatal *Em×1Cre;βl-Integnď°×’°×* animals, n=20 control littermates; Fisher Exact Test p<0.0001). On the other hand, *Ne×Cre;βl-Integrir^flox/flox^* and control P7 cortices are indistinguishable from each other: cortical layers are uniform and defined, as previously shown [45] (**S10A-S10C Fig**). More importantly, no evidence of cortical dysplasia was observed in these mice (**S10 Fig; Table 1;** 0% for both control and *Ne×Cre;βl-Integrir^flox/^f^lox^,* n=10 and n=8 postnatal animals respectively; Fisher Exact Test p=1). This reinforces that *β1-Integrin* acts in an RGC-autonomous manner similar to *Dag1* and *Cas* genes to control cortical lamination.

**Fig. 12.**
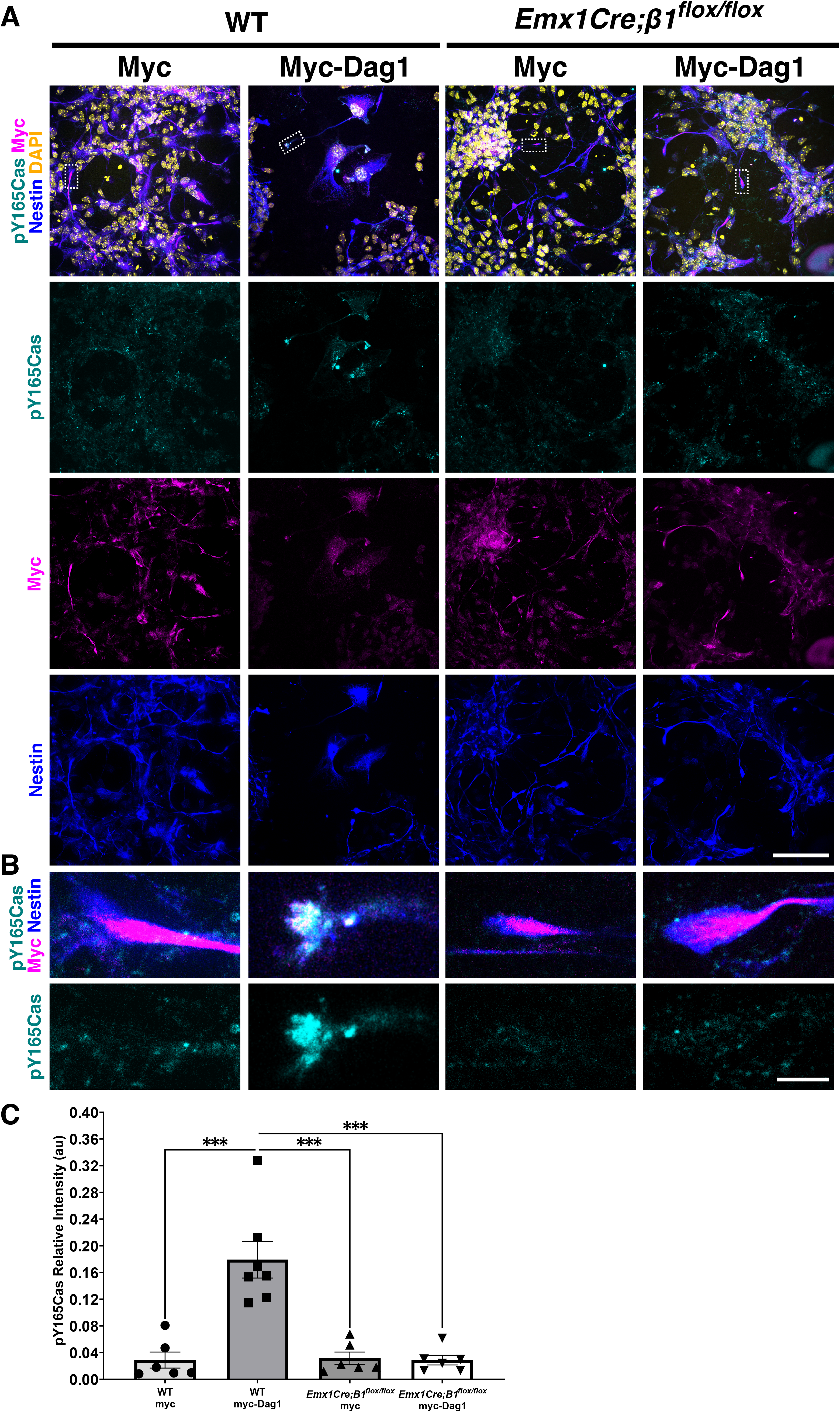
Dag1 and β*1-*Integrin act upstream of Cas phosphorylation in RGC. **(A-B)** Immunofluorescence for pY165Cas (cyan), c-Myc (magenta) and Nestin (blue) of WT and *Em×1Creβ1-Integrin^flox/flox^* mixed cortical cultures transfected with a control vector (myc) or a construct overexpressing myc-tagged full-length Dag1 (myc-Dag). Counterstain in (**A**) is DAPI (yellow). (**B**) is a high magnification view of a representative endbulb boxed in (**A**). Dag1 recruited pY165Cas to the radial glial endbulbs. In the absence of *β1-Integrin,* overexpression of full-length Dag1 does not result in an increase in p130Cas phosphorylation. **(C)** Quantification of the relative fluorescent intensity of pY165Cas at RGC endbulbs. Values given are Mean ± S.E.M., n=6-7 independent samples per group; 9-18 cells per sample, Mann-Whitney *U* test, ***p=0.0001 two-tailed test. For data plotted in graphs, see S13 Data. Scale bars for A: 25 µm; B: 5 µm

**Fig. 13.**
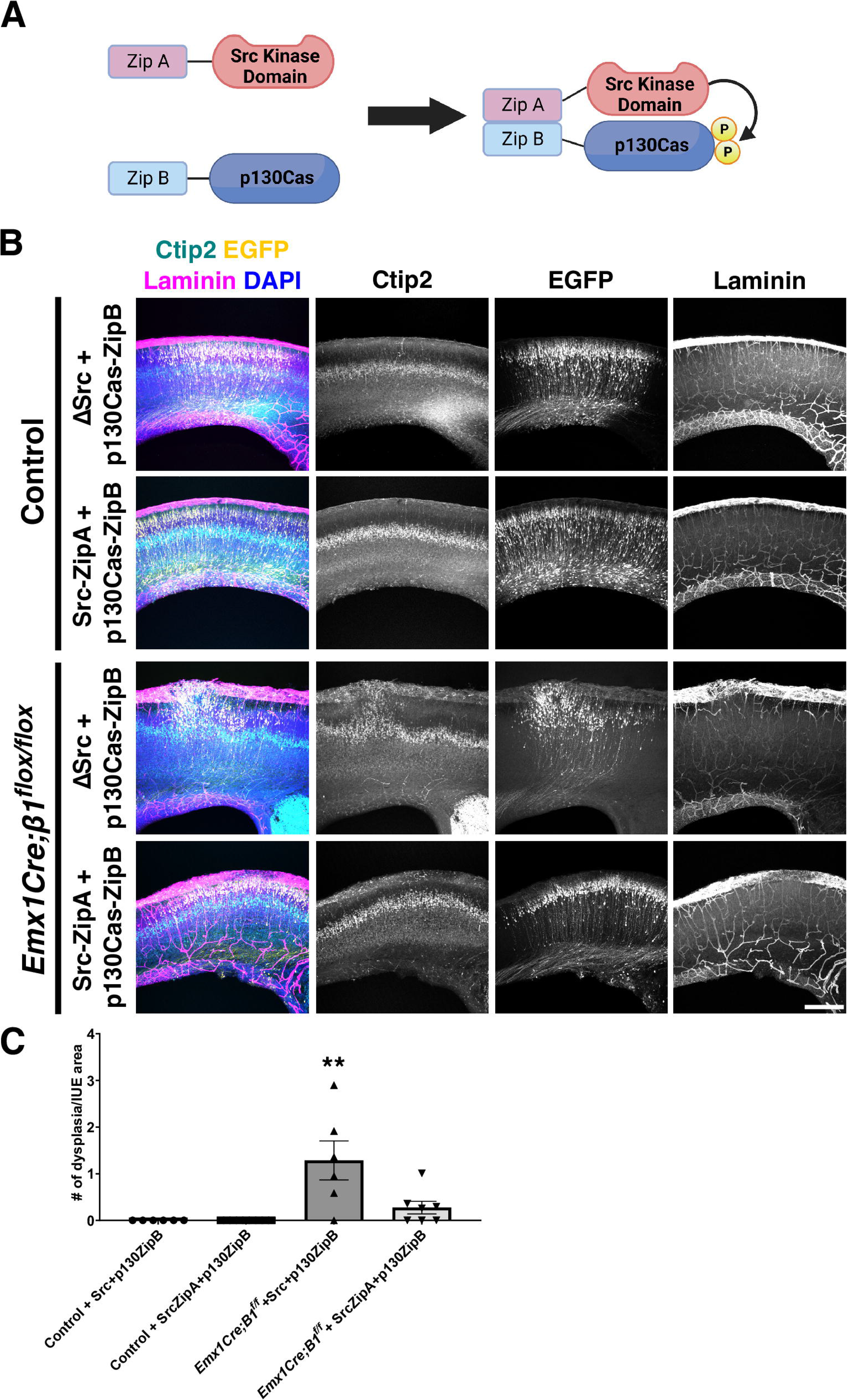
Forced phosphorylation of p130Cas effectively rescues the cobblestone cortex in *β1-Integrin* mutants. **(A)** Schematic representation of Src-ZipA and p130Cas-ZipB constructs used for IUE. The ΔSrc construct is very similar to Src-ZipA, but missing the ZipA domain. (**B**) Coronal cortical sections of Control or *Em×1Cre;βl-Integrin^flox/flox^* P0 animals that were co-electroporated with ΔSrc + p130Cas-ZipB (negative control) or Src-ZipA + p130Cas-ZipB (FIT rescue). These combinations of plasmids were electroporated with pCAGGS-EGFP. Sections were immunostained for EGFP (yellow), Ctip2 (cyan) and laminin (magenta). Nuclei were counterstained with DAPI (blue). Note the evenly layered distribution of Ctip2+ cells in *Em×1Cre;βl-Integrin^flox/flox^* cortices after IUE with Src-ZipA + p130-ZipB. (**C**) Quantification of rescue experiments. Values given are Mean ± S.E.M., n=6-9 independent samples per group; ANOVA p<0.01. Tukey HSD ** p<0.01 for *Emx1Creβ1-Integrin^floxflox^* + ΔSrc + p130Cas-ZipB vs. all other groups. There is no significant difference between *Em×1Cre;βl-Integrin^flox/flox^* + Src-ZipA + p130Cas-ZipB (FIT rescue) and Control animals electroporated with either combination of plasmids. For data plotted in graphs, see S14 Data. Scale bar: 250 µm.

Based on the fact that Dag1 is sufficient to increase pY165Cas in neural cells even in the absence of its cytoplasmic domain, the previously established role for β1-Integrin in regulating Cas function during retina development [61], and the close resemblance between the *Em×1Cre;CasTcKO, Emx1Cre;Dag1^flox/^’*, and *Em×1Cre;βl-Integrin^flox/flox^* cortical phenotypes, we hypothesized that β1-Integrin could be acting as a signal transducing receptor in RGC for this Dag1-dependent increase in Cas phosphorylation. To begin to assess this possibility, we first established whether Dag1 expression can modulate Cas tyrosine phosphorylation in WT RGCs as it did in Neuro2A cells. Mixed primary cultures containing radial glial cells and neurons were transfected at two days *in vitro* with either an empty vector or myc-Dag1. The presence of radial glial cells was confirmed using previously validated developmental markers [119,120] (**S11 Fig**). Subsequently, RGCs were identified by co-labeling with Nestin. Nestin^+^ WT RGCs transfected with an empty vector (pcDNA3.1-myc/his) showed basal endogenous pY165Cas, where puncta are distributed broadly in cell bodies (**Fig 12A).** In Nestin^+^ myc-Dag1-transfected WT cells, we observed a significant accumulation of pY165Cas in the glial endbulbs when compared to control transfected WT cells (**Fig 12A-12C**, Mann-Whitney *U* test, ***p<0.0001 two-tailed test). This finding suggests that Dag1 can modulate p130Cas phosphorylation and may be acting in the same pathway to regulate RGC endbulb interaction with the pial basement membrane.

To test whether Dag1-dependent phosphorylation of Cas proteins requires β1-Integrin as a signal transducing receptor, *Emx1Cre;β1-Integrìn^flox/flox^* cultures were transfected with the empty vector or full-length Dystroglycan (**Fig 12A and 12B**). Analysis using the Kruskal-Wallis test provided very strong evidence of a difference (p=0.002) between the mean ranks of at least one pair of treatments, where a series of Mann-Whitney *U* tests indicated a significant difference between WT RGCs transfected with myc-Dag1 and all of the other groups (**Fig 12C**; ***p<0.001 two-tailed test). Importantly, *Em×1Cre;β1-Integπn^ff¤x/flox^* cells overexpressing myc-Dag1 yielded weak pY165Cas expression in radial glial endbulbs, in comparison to WT cultures overexpressing myc-Dag1 (**Fig 12A-12C**, Mann-Whitney *U* test, ***p<0.001 two-tailed test). Furthermore, no significant difference was observed between *Em×1Cre;βl-Integrin^flox/flox^* RGCs transfected with the control vector or myc-Dag1 (**Fig 12A-C)**. These data indicate that Dagl-dependent phosphorylation or recruitment of pYCas to the endbulbs requires β1-Integrin function.

These tissue culture experiments and the resemblance between the cortical phenotypes of the *Dag1, β1-Integrin* and *Cas* pan-cortical knock-outs, while suggestive of a possible interaction, are in no way definitive proof of an epistatic relationship between these genes. To directly test whether Cas adaptor proteins act downstream of β1-Integrin to regulate cortical lamination we performed a rescue experiment. We reasoned that if β1-Integrin-dependent signaling is critically mediated by Cas protein phosphorylation to regulate cortical migration and lamination, forcing p130Cas tyrosine phosphorylation into RGC might be able to rescue a β1-Integrin deficiency. With this in mind, we took advantage of the functional interaction trap (FIT) system [121,122]. FIT constructs allow for the tethering of a kinase to its target through an engineered, highly specific binding interface, using a pair of complementary synthetic amphipathic helices (coiled coils) [121,122]. This highly specific binding interface is used to replace their natural interaction sites, and results in efficient phosphorylation of the substrate. To test whether forced p130Cas phosphorylation could rescue the formation of cortical dysplasia in *Emx1Cre;β1-Integrin^flox/flox^,* we co-electroporated FIT constructs that prompt the interaction between Src and Cas into the cortices of these embryos and littermate controls at E14.5 and collected them at P0. For Src, we used a construct lacking the substrate binding SH2 and SH3 domains, but containing a coiled-coiled domain (ZipA) and an intact membrane targeting sequence (Src-ZipA) (**Fig 13A**). The target p130Cas was fused to ZipB, the coiled-coil segment complementary to ZipA (p130Cas-ZipB) (**Fig 13A**) [121,122]. Due to the difficulty of these experiments and the possibility of the plasmid titer diluting out inside RGCs as they divide, we opted for driving the expression of these constructs with a strong constitutive promoter *(EF-1α)* rather than weaker RGC-specific ones. As negative control, we co-electroporated a similar deletion construct for Src lacking the ZipA coiled-coil domain (ΔSrc) and p130Cas-ZipB. These combinations of constructs were co-electroporated with pCAGGS-EGFP to visualize the electroporated area. Coronal sections of P0 electroporated brains were then stained for Laminin to label the basement membrane, and Ctip2 to visualize the cortical lamination pattern. Co-electroporation of the negative control constructs (ΔSrc + p130Cas-ZipB) into *Emx1Cre;β1-Integnn^flox/+^* or *β1-Integrin^flox/flox^* control animals did not affect cortical lamination or basement membrane stability (**Fig 13B and 13C**). Importantly, neither did co-electroporation of Src-ZipA + p130Cas-ZipB into the same control animals (**Fig 13B**). When ΔSrc + p130Cas-ZipB were electroporated into *Emx1Creβ1-Integrin^flox/flox^* the characteristic cobblestone cortex with obvert dysplasia was observed, even in the EGFP+ region (**Fig 13B and 13C**). This was comparable to the presence of dysplasia on the uninjected contralateral side (**S12 Fig**). However, introducing Src-ZipA + p130Cas-ZipB into the *Em×1Creβ1-Integrin^flox/flox^* mutant cortices notably rescued the cobblestone phenotype within the EGFP+ electroporated region (**Fig 13B and 13C**): Ctip2+ cells are positioned in their normal laminar location, with little to no evidence of dysplasia or ectopias breaking the basement membrane (One-Way ANOVA, p=0.0003 two-tailed test; Tukey HSD **p<0.01 Src + p130Cas-ZipB IUE into *Em×1Cie;βl-Integrin^flox/flox^* vs. Src-ZipA + p130Cas-ZipB IUE into *Em×1Creβ1-Integrin^flox/flox^;* **p<0.01 Src + p130Cas-ZipB IUE into *Em×1Cre;β1-Integrin^flox/flox^* vs. both IUEs into control animals; *ns:* Src-ZipA + p130Cas-ZipB IUE into *Emx1Cre;β1-Integrin^flox/flox^* vs. both control lUEs; 6-9 animals per treatment per genotype). This rescue was specific to the electroporated region since the contralateral side still displayed noticeable dysplasia (**S12 Fig**). These results unequivocally place Cas protein tyrosine phosphorylation downstream of β1-Integrin during cortical lamination. Furthermore, the combination of these genetic and tissue culture experiments strongly suggest that Dag1, β1-Integrin and Cas act in the same pathway and in a non-neuronal autonomous manner to regulate basement membrane integrity and cortical lamination.

## Discussion

Lamination and stratification of neuronal cell bodies are widespread early developmental events that organize many regions of the CNS [5,12,13,16,123,124]. These foundational processes are thought to provide a structure upon which neural circuits can form [125,126,127]. During many of these stratification events, migrating neurons utilize RGCs to reach the proper layer [10,128,129,130]. In this study we describe a molecular mechanism by which the glial scaffold is assembled to facilitate cortical migration and lamination. Our data identify an adhesion signaling axis that acts in RGCs to maintain glial endfeet-pial interactions and establish a novel and essential role for Cas proteins in these developmental processes.

Assembly and maintenance of the glial scaffold is particularly important during neocortical lamination [10,47,131]. The formation of this scaffold requires proper attachment of the glial endfeet to the pial basement membrane. Disruption of these interactions in mouse and human results in severe lamination defects like cortical dysplasia and cobblestone cortex [46,47,50,132]. Why is constant engagement of the basal lamina by RGCs so critical? While the cortical scaffold is traditionally portrayed as a somewhat rigid structure, the cortex expands vastly during neurogenesis [133,134,135]. RGC processes must grow and remodel to accommodate this rapid expansion of the cortical plate as new layers are born and incorporated [102,131]. RGC endfeet must actively adhere to the basement membrane to maintain the structure of the scaffold [40,131,136]. Thus, dynamic regulation of adhesion signaling and cytoskeletal remodeling are essential for these glial-pial interactions.

Our data support a role for Cas adaptor proteins acting as signal transducers downstream of the β1-Integrin and Dystroglycan transmembrane proteins to regulate RGC-basement membrane interactions and maintain the integrity of the glial scaffold. *In vitro* data further suggests that the Dag1-dependent phosphorylation of Cas proteins is mediated by β1-Integrin (**Fig 10C** and **Fig 12**) and data from rescue experiments reinforce this idea (**Fig 13**). This signaling axis is likely modulated by other transducers of Integrin signaling known to act in non-neuronal-autonomous manner to establish or maintain the RG scaffold [8,137,138]. While there is a clear disruption of the glial scaffold in the *Em×1Cre;CasTcKO* animals, the alterations to the dorsal and ventral boundaries of the CP and clustering of Cajal-Retzius cells observed in these animals also likely contribute to the lamination defects [36,97,139,140,141], similar to what occurs in *Dag1* mutant cortices [46,47,142,143,144]. Based on phenotypic analyses, genetic rescue, and molecular epistasis data we propose a working model whereby the Dag1-Integrin-Cas adhesion axis acts in at least two ways (**Fig 14**). In cis, it allows RGCs to anchor and remodel their endfeet as the cortex grows, promoting the establishment and maintenance of the glial scaffold by interacting with the ECM. Dag1 and Integrin in the RGC endfeet also act in trans through their extracellular domains to organize and stabilize the basement membrane by nucleating or interacting with ECM molecules like Laminin. Proper establishment and maintenance of the glial scaffold and basement membrane allow for normal neuronal migration and lamination. While this model is supported by genetic experiments by others and us [8,43,45,46,47,137,138,145], further studies will be needed to test this at the molecular level and to probe whether other genes with similar roles in glial scaffold formation act in parallel or in the same signaling pathway [168, 169].

**Fig. 14.**
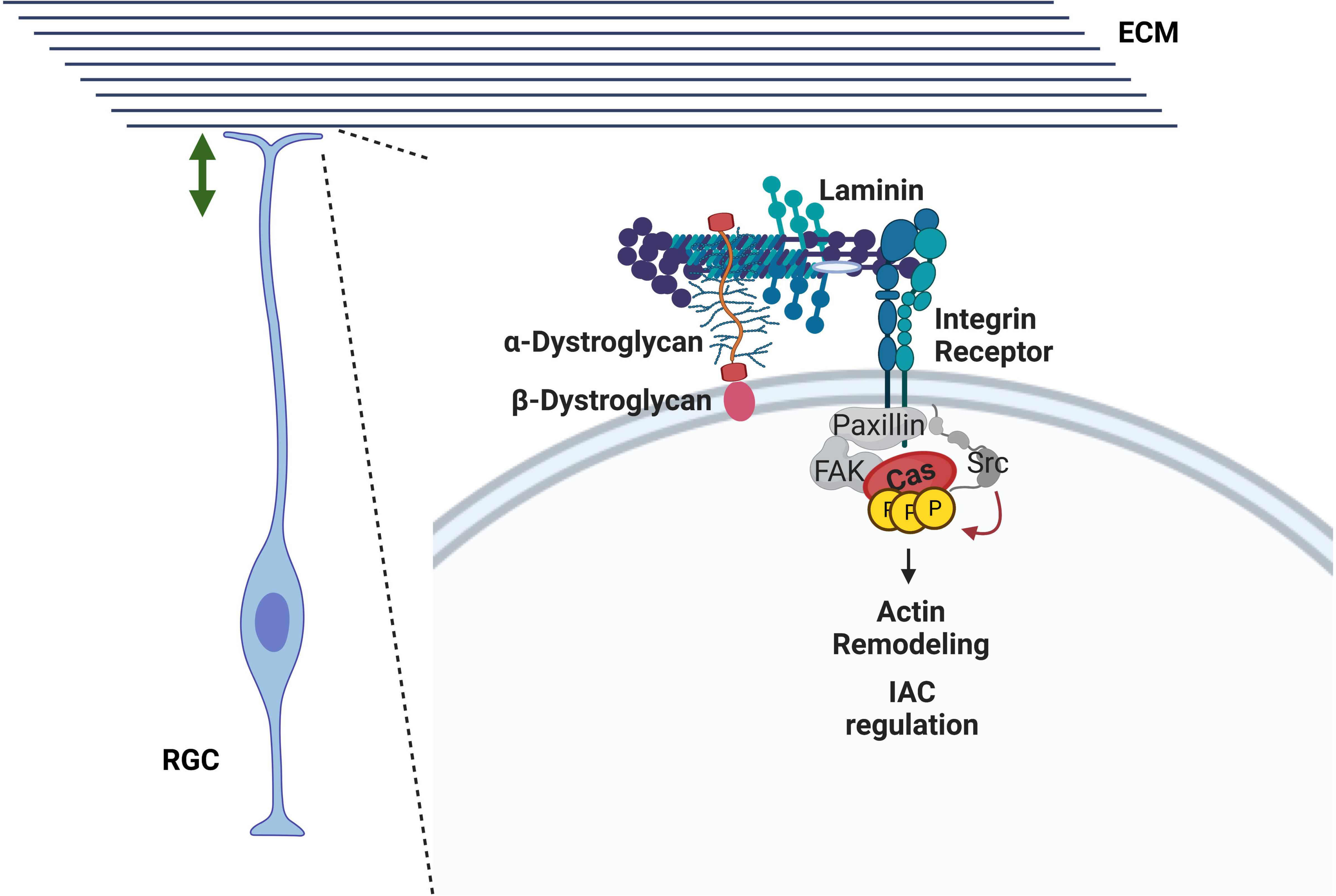
A working model for an adhesion signaling axis regulating RGC scaffold formation and basement membrane maintenance. A Dag1-Integrin-Cas axis acts in *cis* in RGCs to maintain attachment to the basement membrane as the cortex expands. In addition, Dag1 and β1-Integrin participate in the maintenance of the basement membrane by acting in *trans* through their extracellular domains. Additional cytoplasmic IAC proteins that appear in grey have been showed to participate in cortical lamination based on genetic evidence and we speculate that they could be acting in the same pathway [29,30,31,32,33,34,35].

The findings presented here add to a growing list of studies in a variety of animal models that have demonstrated the evolutionarily conserved requirement for Cas adaptor proteins during cell migration and axon pathfinding [59,60,61,62]. In the cortex, the aforementioned Integrin-Cas axis appears to act primarily in a glial-autonomous manner [43,45]. Genetic evidence and gene expression data strongly support this function of Cas family members in RGC. However, since we were unable to detect specific phospho-Cas signal *in vivo,* it is still possible that Cas proteins influence endfeet attachment and glial scaffold organization by acting in other subcellular compartments of RGC. Contrary to its glial-autonomous function in the cortex, previous work demonstrated a neuronal-autonomous function for a similar signaling pathway in mouse and fly peripheral axon pathfinding, and mammalian retina development [59,61,62]. During retina lamination and stratification, neurons initiate migration without the aid of radial glia and the Laminin-rich basement membrane provides an instructive cue that defines the position where the single-cell ganglion cell layer will form [61,146,147,148,149,150]. In the eye the Integrin-Cas signaling axis acts in migrating retinal ganglion cells to sense the inner limiting membrane and is necessary for the reorganization of the ganglion cell layer [61]. Of note, even though Crk-Associated Substrates (Cas) are essential for RGC process and endfeet organization, histological and biochemical evidence support the idea that the Crk proteins themselves function in a neuronal-autonomous manner downstream of reelin signaling to regulate cortical lamination [41,90,151].

Cobblestone or type II lissencephaly is a debilitating neurodevelopmental disorder caused by breaks in the basement membrane (BM) at the pial surface [50,152,153,154]. These defects have been associated with RGC endfeet that are not well attached to the ECM [131,155]. The BM breakages result in the over-migration of neurons and a dramatic loss of normal cortical lamination [37,46,47,142,156]. The genetic causes of cobblestone lissencephaly have largely been linked to autosomal recessive mutations in genes that encode for proteins that participate in the post­translational modification of alpha-Dystroglycan: POMT1, POMT2, POMGNT1, FCMD/FKTN, FKRP, TMEM5/RXYLT1, ISPD/CRPPA and LARGE1 [46,50,107,157,158]. Mutations in one of these genes are found in 32–66% of patients with Type II lissencephaly [50,158,159,160].However, despite intensive research in this area, more than one-third of the cases have an unknown genetic etiology [50].The striking resemblance of the phenotypes observed in cortical *CasTcKOs* and patients with cobblestone lissencephaly, in addition to the molecular epistasis experiments suggesting that Cas proteins act downstream of Dag1 and β1-Integrin in radial glial cells to maintain basement membrane integrity, pinpoint cytoplasmic effectors of adhesion signaling as possible contributing factors to the etiology of these neurodevelopmental disorders. A recent study revealed that several genes involved in regulating cell-matrix adhesion and IAC assembly are somatically mutated in patients with Focal Cortical Dysplasia [161]. Patients with FCD type I show disruptions of the cortical laminar structure that parallel those observed in Type II lissencephaly, but as the name implies, these dysplasia are more focal in nature, i.e. restricted to a small region of the cortex [154,162]. Putting our study in the context of these data, our findings open new and intriguing avenues in the study of the genetic causes of cobblestone lissencephaly and possibly FCD.

## Materials & Methods

### Animals

The day of vaginal plug observation was designated as embryonic day 0.5 (E0.5) and the day of birth was postnatal day 0 (P0). The ICR mouse strain was purchased from Taconic. The following animals were purchased from The Jackson Laboratory: *Ai14/R26^LSL-TdTomato^* (stock 007914), *β1-Integr^f^* (stock 004605), *Em×1Cre* (stock 005628), and *Dag1^flox^* (stock 009652). *NexCre* mice were kindly provided by Drs. Nave and Goebbels [99,100]. Generation of the *p130Cas^flox/flox^, CasL^-/-^* and *Sin^-/-^* transgenic mouse lines has been described previously [61,68,69]. The *Dag1^βcyto^* mice were kindly provided by Dr. Kevin Campbell and previously described in [112]. Generation and characterization of *p130Cas::BacEGFP* animals was described in [61,62, 66, 67]. All animal procedures were performed according to the University of California, Riverside’s Institutional Animal Care and Use Committee (IACUC) guidelines.

### Immunofluorescence

Embryonic brains were dissected in cold PBS and postnatal mice were perfused then dissected. The brain samples were post-fixed in 4% PFA for 2 Hrs., 4 Hrs., or overnight at 4°C and washed with PBS the next day. The samples were either kept in PBS or sunk in 30% sucrose overnight at 4°C prior to cryopreservation with Optimum Cutting Temperature (Tissue-Tek). Brains were either embedded in 3% agarose for coronal sectioning at 150 µm thickness on a vibrating microtome (VT1000S; Leica) or cryoprotected samples and sectioned coronally at 20 µm on a Leica CM3050 cryostat. Cryosections were blocked for 1 hour in RT in blocking buffer (0.1% PBST and 10% goat serum) in a humidified chamber and free-floating slices were blocked in permeabilization solution overnight at 4°C as previously described [163]. Primary antibodies were diluted in antibody dilution buffer (cryosections; 0.1% PBST, 1% goat serum) or permeabilization solution (free-floating slices) with the following antibodies: rabbit anti-Cux1 (1:250, ProteinTech# 11733-1-AP), rat anti-Ctip2 (1:500, Abcam# ab18465), rabbit anti-Tbr1 (1:500, Abcam# ab31940), rabbit anto-Tbr2 (1:500, Abcam# ab23345), rabbit anti-Rorβ (1:250, Proteintech# 17635-1-AP), rabbit anti-Calretinin (1:500, Swant# 7697), rabbit anti-Calbindin (1:500, Swant# CB38), chicken anti-MAP2 (1:500, Aves MAP), rat anti-NCAM-L1 (1:500, Millipore# MAB5272), rabbit anti FoxP2 (1:500, Abcam# ab16046), chicken anti-EGFP (1:1000, Aves Labs# GFP-1020), rabbit anti-EGFP (1:1000, Thermo Fisher# A-6455) and rabbit anti-CSPG (1:1000; Sigma# AB5320). The samples incubated in the primary antibody mix overnight at 4°C. Cryosections were washed with PBS 15 mins × 4 in a slide mailer and incubated in secondary antibody (goat anti-chicken 647, Themo Fisher Scientific Cat# A-21449 and goat anti-rabbit 546, Thermo Fisher Scientific Cat# A-11035) with DAPI (1 µg/ml) and antibody dilution buffer at RT for 1 hour in a humidified chamber. Slides were washed with PBS 15 mins × 4 in a slide mailer and sealed with fluorogel. Free floating sections were washed with PBS 15 mins × 8 and incubated in secondary antibody at 1:1000 dilution in permeabilization solution overnight at 4°C. Sections were washed with PBS 15 mins × 4 and sealed with fluorogel. Samples were imaged with a confocal microscope.

### Ethynyl deoxyuridine (EdU) Labeling

EdU was dissolved in DMSO and diluted with sterile water to 15 mg/ml at the time of injection. EdU was injected into the intraperitoneal cavity of pregnant mice at a dosage of 150 mg/kg. Animals were collected 30 minutes or 7 days after administration. Embryonic brains were dissected in cold PBS and postnatal animals were perfused then dissected. The brains were fixed for 2 hours in 4% PFA, dehydrated in 30% sucrose overnight, then cryopreserved with Optimum Cutting Temperature (Tissue-Tek). Cryosections were stained according to the instructions in the Click-iT EdU Cell Proliferation Kit for Imaging, Alexa Fluor 488 dye, Invitrogen Cat# C10337. Slides were incubated with 3% BSA for 30 mins at RT, then blocked in 0.5% PBST at RT for 1 hour, washed with 3% BSA in PBS × 2, and incubated in Click-iT Reaction Cocktail (1x Click-iT Reaction Buffer, CuSθ_4_, Alexa Fluor Azide 488, and Reaction Buffer Additive) for 1 hour at RT. Cryosections were washed with 3% BSA in PBS × 4 and immunohistochemistry proceeded as described [163]. Primary antibodies chicken anti-Nestin (1:500, Aves Nes) and rabbit anti-Laminin (1:500, Sigma# L9393) were diluted in antibody dilution buffer (0.1% PBST, 1% goat serum). Slides incubated in the primary antibody mix in a humidified chamber at 4°C overnight. Slides were washed with PBS 15 mins × 4 in a slide mailer and incubated in secondary antibody at 1:1000 dilution (goat anti-chicken 647, Themo Fisher Scientific Cat# A-21449 and goat anti-rabbit 546, Thermo Fisher Scientific Cat# A-11035) at RT for 1 hour in a humidified chamber. Slides were washed with PBS 15 mins × 4 in a slide mailer. Fluorogel was applied to the slide and sealed with a coverslip. Samples were imaged with a confocal microscope.

### Mixed Primary Cell Culture

Primary mixed radial glial cells and cortical neurons were isolated from E13.5 embryos essentially as described [164] using Complete Hank’s Balanced Salt Solution (HBSS, 2.5 mM HEPES pH 7.4, 2.5 mM D-glucose, 1 mM CaCl_2_, 1 mM MgSO_4_, 2.5 mM NaHCO_3_) [163]. Cells were dissociated as described [165] and grown in Radial Glia Media modified from [165] (Modified DMEM/Eagle’s Media, 1x Penicillin/Streptomycin, 0.5 mM Glutamax, 20 mM D-glucose, and 5% horse serum). 12 mm circular coverslips were treated with 12 M HCL overnight, neutralized with deionized water, and stored in a glass petri dish with 70-100% ethanol. The coverslips were fire-polished into a 24-well plate and coated with Laminin (8.3 µg/ml)) and poly-D-lysine/poly-L-lysine (83.3 µg/ml) [163] in sterile water overnight in a tissue culture incubator at 37°C in 5% CO_2_. The coating media was aspirated and the coverslips were air-dried in an airflow cabinet prior to plating cells. Dissociated cells were plated in a 24-well plate at a density of 1 × 10^5^ cells/well with 500 µl of cell suspension and Radial Glia Media. One-third of the media was replaced daily and wells were transfected two days *in vitro* (2DIV) with Metafectene PRO (Biontex Cat# T040-1.0), according to the manufacturer’s suggested protocol of 1 µg DNA to 2 µl Metafectene ratio. Cells were transfected with either pcDNA3.1-myc/his (control vector) or pcDNA3.1-myc/his-Dag1-FL (myc-Dag1) plasmids previously described [166]. At 48 hours after transfection (4DIV), the coverslips were fixed in 4% PFA for 10 minutes at RT and washed with PBS with 0.03% NaN3. Cells incubated in blocking buffer (0.1% PBST, 10% goat serum) for 30 minutes at RT. Un-transfected cells were stained with the following primary antibodies: rabbit anti-Blbp (1:500, Abcam# ab32423), mouse anti-β3-Tubulin (1:500, CST# 4466), rat anti-GFAP (1:500, Invitrogen# 13-0300), rabbit anti-Tbr2 (1:500, Abcam# ab23345), and chicken anti-Nestin (1:500, Aves Nes) in antibody dilution buffer (0.1% PBST, 1% goat serum) overnight at 4°C. Transfected cells incubated in primary antibodies: mouse anti-c-myc 9E10 (1:1000, Invitrogen# 13-2500), rabbit anti-Tyr165 (1:250, CST# 4015), and chicken anti-Nestin (1:500, Aves Nes) in antibody dilution buffer overnight at 4°C. Cells were then washed with PBS with 0.03% NaN_3_ for 15 mins RT × 4 and incubated in secondary antibodies at 1:1000 dilution (goat anti-mouse 488, Thermo Fisher Scientific# A-28175, goat anti-rabbit 647, Thermo Fisher Scientific# A-27040, goat anti-chicken 546, Thermo Fisher Scientific# A-11040, goat anti-rat 647, Thermo Fisher Scientific# A-21247) and DAPI (1 µg/ml) in antibody dilution buffer at RT for 2 hours. Cells were washed with PBS with 0.03% NaN_3_ for 15 mins × 4 and coverslips were mounted on microscope slides with fluorogel and imaged with a Leica confocal microscope.

#### Neuro2A Cell Culture

Neuro2A cells were obtained from ATCC and incubated at 5% CO_2_ at 37°C. The growth media was DMEM with pen/strep (Gibco Cat#15140-122) supplemented with 10% fetal bovine serum. Cells were visually inspected for contamination daily and routinely tested for Mycoplasma infection using ATCC Universal Mycoplasma Detection Kit (ATCC# 30-1012K). Cells were plated on Laminin and poly-D-lysine (8.3µg/ml) coated 12 mm coverslips in a 24-well plate at 7.8×10^4^ cells/cm^2^. Cells were transfected 24 hours later with Metafectene PRO (Biontex Cat# T040-1.0) with according to the manufacturer’s suggested protocol with a 1:4 DNA to Metafectene ratio. Cells were transfected with either pcDNA3.1-myc/his (control vector) or pcDNA3.1-myc/his-Dag1-FL (myc-Dag1) or pcDNA3.1-myc/his-Dag1-FC (myc-Dag1_ecto_) plasmids previously described [166]. Cells were fixed 48 hours after transfection with 4% PFA for 10 minutes at RT, rinsed with PBS, and immunostained as described above for primary cultures.

### Quantification of Layer Markers, EdU and ectopias, and Statistical Analysis

Cells were counted using the RapID Cell Counter [167]. The thickness of the cortex was divided into 10 bins of equal width and the number of cells labeled with the layer markers (Cux1, Ctip2, Rorβ, Tbr1) and/or EdU were counted for each bin. Bin 1 is the marginal zone (MZ) and bin 10 is ventral to layer VI. The proportion of cells in each bin= number of cells labeled in bin divided by total number of labeled cells. The graph values are mean ± standard error of mean, n=3-5 independent animals per group depending on the experiment, and on average 3 sections per sample. For experiments quantifying density, area or length were quantified using ImageJ. Cell counts for neurons labeled by one or more markers were performed using RapID Cell Counter [167]. Number of dysplasia or ectopias were counted manually using the ImageJ multi-point tool. The datasets were then tested for normality using the Shapiro-Wilk test and QQ plot. For normally distributed datasets, two tailed t-test was performed for comparing two samples, and one-way ANOVA followed by Tukey HSD was used for multiple comparisons. For datasets that didn’t fit a normal distribution the Mann-Whitney *U* test was performed with Bonferroni correction (e.g. p<0.005 was required for significance for binning experiments with 10 pairwise comparisons).

### Mean Fluorescent Intensity of immunofluorescence

ImageJ (NIH) was used to measure the fluorescent intensity of anti-Nestin and anti-Tyr165 in radial glial endbulb. The Nestin channel was used to trace the borders for the radial glial endbulb. The following selections were used to determine the fluorescent intensity of Nestin and pY165Cas channels using ImageJ: Analyze -> Set Measurements (Area, Min & max gray value, Integrated density, Mean gray value)-> Measure mean gray value. Relative fluorescent intensity is mean fluorescent intensity of pY165Cas divided by mean fluorescent intensity of Nestin. The graph values are mean ± standard error of mean, n=6-7 independent samples per group, and 9-18 cells per sample. The dataset was tested for normality using the Shapiro-Wilk test and QQ plot. For the mixed cultures Kruskal-Wallis test was performed for multiple-comparisons, followed by a series of Mann-Whitney *U* tests to examine which pairs are significantly different. For Neuro2A cells, pY165Cas signal was normalized to transfected cells on the same image, since expression of myc in the absence of a fusion protein is very low. For these experiments, datasets were normally distributed and thus ANOVA followed by Tukey HSD test was performed.

#### *In utero* electroporation

Pregnant female mice were anesthetized with isoflurane. An abdominal incision was made to expose the uterus and intrauterine electroporation was then conducted as described in [170]. Briefly, pups were visualized through the uterine wall. Plasmids diluted in fast green and sterile PBS were injected through sharpened glass capillary needles into the lateral ventricle. 5mm paddles were used to deliver five 35V (E13.5) or 37V (E14.5) pulses of 50 ms each with 950 ms intervals across the hemispheres. After electroporation, uterine horns were repositioned inside the female and the abdominal cavity was filled with warm 5x penicillin/streptomycin (pen/strep) in sterile PBS. For labeling with EGFP under the *BLBP* promoter, E13.5 WT ICR embryos were injected and electroporated with 3ug pBLBP-EGFP. Pups were collected at E16.5 and processed for RNAscope as described above. For rescue experiments, E14.5 embryos from a *β1-Integrin^flox/flox^* dam crossed to an *Em×1Cre;β1-Integnn^f¤xX+^* sire were electroporated with a 1:1:1 ratio ΔSrc + p130Cas-ZipB + pCAGGS-EGFP mix or Src-ZipA + p130Cas-ZipB + pCAGGS-EGFP using 2 µg of each plasmid/IUE. Brains were collected at P0 after transcardial perfusion with PBS and 4% paraformaldehyde (PFA), and fixed in 4% PFA for four hours at 4°C. Brain we then rinsed with PBS, and sectioned coronally at 150 µm on a vibrating microtome (VT100S; Leica) and processed for immunofluorescence. BLBP-GFP was a gift from Bart De Strooper (Addgene plasmid # 63174).

### Fluorescent in situ Hybridization

Fluorescent *in situ* hybridization was performed on 20 µm E12.5, E14.5 or E16.5 cryosections using the RNAScope Multiplex Fluorescent Detection Kit v2 from Advanced Cell Diagnostics according to manufacturer instructions (ACD, 323110) [65]. Briefly, target RNAs are hybridized to ssDNA “z-probes” complementary to RNA of interest. Oligos bind to the tail region of the z-probe, which are then bound to amplifiers labeled with horse radish peroxidase (HRP) and fluorophores. *p130Cas/BCAR1* (ACD, 806961), *CasL/NEDD9* (ACD, 446311), and *Sin/EFS* (ACD, 806951) probes generated by Advanced Cell Diagnostics were uniquely amplified with Opal dyes (Akoya Biosciences, FP1487001KT, FP1488001KT, FP1496001KT) and then counterstained with DAPI (1 µg/ml). Slides were mounted using Fluoro-Gel mounting medium (Electron Microscopy Sciences, 1798510) and imaged on a Leica SPE confocal microscope. Quantification was performed in ImageJ: the different cortical areas were outlined based on the DAPI channel, and the fluorescent channel corresponding to the RNA was then made binary. The watershed function was then applied, and particles were analyzed within the ROI using the “analyze particles” function. All counts are reported with ± standard error of the mean [171].

## Acknowledgements

We would like to thank Drs. Sachiko Haga-Yamanaka and Edward Zagha for critically reading the manuscript and providing helpful comments. We would like to thank Drs. Nave and Goebbels for the *NexCre* mice. We would also like to thank Drs. Sachiko Seo and Mineo Kurokawa, and Dr. Konstantina Alexandropoulos for sharing the *CasL^-^’* and *Sin^-^’* mouse lines, respectively. We also would like to thank Bruce Mayer for the FIT constructs and Bart De Strooper for the Blbp-EGFP plasmid.

## Availability of data and materials

All data analyzed during this study are included in this article. The data underlying all the graphs is presented in **S1 Data**-**S14 Data.**

## Authors Contributions

WW and JNJ designed and performed experiments, analyzed data and wrote the manuscript. AMT, NR, JAE, TU, and KMW designed and performed experiments, and analyzed data. JAE and KMW also provided comments and suggestions for the manuscript. MMR conceived the project, designed experiments, analyzed data and wrote the manuscript.

## Ethics approval

All animal procedures presented here were performed according to the University of California, Riverside’s Institutional Animal Care and Use Committee (IACUC)-approved guidelines, under protocol A-20200021

## Supplemental Figure Legends

**S1 Fig. EGFP antibodies are specific for EGFP. (A-B)** Immunostaining of cortical sections with the same chicken-anti EGFP (**A**) and rabbit-anti EGFP (**B**) antibodies used to detect it in *p130Cas::BacEGFP* animals produce negligible signal on WT sections at E12.5 and E15.5. Magenta channel in (**A**) is Tbr1 and in (**B**) is Nestin. Dapi counterstain is shown in blue. n=3 animals. PP: preplate, CP: cortical plate; IZ: intermediate zone; SVZ: subventricular zone; VZ: ventricular zone. Scale bars for A and B (top): 25 µm; A and B (bottom): 75 µm.

**S2 Fig. *Em×1Cre* efficiently recombines in the developing cortex.** (**A**) tdTomato staining of E13.5 *Emx1Cre;Ai14* coronal cortical sections. *Em×1Cre* drives Cre recombinase expression in cortical neural progenitors and radial glial cells that generate excitatory neurons in the forebrain and hippocampus. (**B**) RNAscope at E14.5 shows that *Em×1Cre;CasTcKO* cortices do not produce functional *p130Cas* transcripts. Control littermates express *p130Cas* transcripts strongly in the IZ and broadly throughout the neocortex. Neither controls, nor *Em×1Cre;CasTcKO* animals express *Sin* or *CasL.* n=3 animals per genotype. CP: cortical plate; IZ: intermediate zone; SVZ: subventricular zone; VZ: ventricular zone. Scale bars for A: 500 µm; D: 100 µm.

**S3 Fig. Characterization of cortical phenotypes in *Em×1Cre;CasTcKO* mice. (A-C)** Immunostaining of P7 control and *Em×1Cre;CasTcKO* coronal sections using the layer markers Cux1 (green, layer II/III) and Ctip2 (magenta, layer V & VI) (**A**), Rorβ+ (**B**, red, layer IV) and Tbr1+ (**C**, red, layer VI). Neurons appear misplaced in *Em×1Cre;CasTcKO* mice (white arrows). DAPI (blue) was used to counterstain. (**A**) Right panels: quantification of Ctip+ and Cux1+ neuron density. two tailed t-test *ns: p=0.12* for Ctip2+ cells; p=0.09 for Cux1+ cells; n=5 independent animals for each genotype, 3 section per animal. (**B, C**) Right panels: quantification of the proportion of Rorβ+ and Tbr1+ cells per bin. Bin 1 is the MZ and bin 10 is ventral to layer VI. n=3 animals per group, 3 sections per animal, Mann-Whitney *U* test & Bonferroni correction, *p<0.005. Values given are Mean ± S.E.M. For data plotted in graphs, see S2 Data. Scale bars: 500 µm.

**S4 Fig. Analysis of for *CasL, Sin* and *p130Cas* null allelic series.** Coronal sections of P7 *Emx1Cre;p130Cas^flox/+^;CasL^+/-^;Sin^+/-^*, *Em×1Cre; p130Cas^flox/+^;CasL^-/-^;Sin^-/-^*, *Em×1Cre; p130Cas^fo>x-^;CasL+^/-^;Sin^-/-^*, *Em×1Cre; p130Cas^flox/-^;CasL^-/-^;Sin^+/-^* and *Em×1Cre; p130Cas^flox/-^;CasL^-/-^;Sin^-/-^ (Em×1Cre;CasTcKO)* cortices stained for the layer markers Ctip2 (green, layer V & VI) and Tbr1 (red, Layer VI), and the mature neuron marker NeuN (grey), counterstained with DAPI (blue). Only *Em×1Cre; p130Cas^flox/-^ ;Casư^//^;Sın^+//^* and *Em×1Cre; p130Cas^flox/-^;CasL^-/-^;Sin^-/-^* showed the cobblestone cortex phenotype. Bottom Right panel: quantification of the # of dysplasia/mm^2^ in the different transgenic animals. Values given are Mean ± S.E.M., n=3-8 independent samples per group, 3 sections per sample, Mann-Whitney *U* test & Bonferroni correction, *p<0.005 vs. control; ** p<0.0005 vs. control. For data plotted in graphs, see S3 Data. Scale bar: 1 mm.

**S5 Fig. Defective preplate split in *Em×1Cre;CasTcKO* mice.** Immunostaining of control and *Emx1Cre;CasTcKO* coronal sections for marginal zone (MZ) and subplate (SP; white arrows) markers at E15.5. MAP2, CSPG, Calretinin, and Calbindin show clustering of cells (yellow arrows). CSPG+ cells form a column connecting the SP with the MZ (yellow arrow). Thalamocortical afferents labeled with Calretinin show ectopic bundles extending from the intermediate zone (IZ) (white arrowheads). DAPI was used as counterstain (blue). n=3 animals per genotype. Scale bars: 100 µm.

**S6 Fig. Caspase3 activation is not affected in *Em×1Cre;CasTcKO* mice at key developmental stages. (A-D)** Immunostaining for activated Caspase3 (aCasp3, green) on E12.5 (**A**), E15.5 (**B**), P0 (**C**) or P3 (**D**) coronal sections of Control and *Em×1Cre;CasTcKO* cortices. Sections were counterstained with DAPI (blue). No notable differences are observed between genotypes. Right panels: quantification of density of activated Caspase 3. Values given are Mean ± S.E.M., n=3 independent samples per group, 3-5 sections per sample, Mann-Whitney *U* test & Bonferroni correction, ns for all stages (p≥0.7). For data plotted in graphs, see S6 Data. Scale bars for A: 100 µm; B, C, D:500 µm.

**S7 Fig. *NexCre* efficiently ablates *p130Cas^fí¤x^*in postmitotic cortical neurons. (A)** tdTomato staining of E13.5 *NexCre;Ai14* coronal cortical sections. *NexCre* targets postmitotic, premigratory neurons in the neocortex. (**B**) RNAscope at E16.5 indicates that *NexCre;CasTcKO* cortices express negligible levels of functional *p130Cas* transcripts in the IZ and CP, and display significantly less transcript in the VZ and SVZ. Top Right panel: quantification puncta per micron DAPI area in cortical plate + intermediate zone (CP+IZ) was 0.114±0.009 for *Nex-Cre^+^;CasTcKO^flox/+^* and 0.116±0.005 for ventricular + subventricular zones (VZ+SVZ). For *Nex-Cre^+^;CasTcKO^flox/^’’* the average *p130Cas* puncta per micron DAPI area in CP+IZ was 0.013±0.003 and 0.056±001 for VZ and SVZ. Values given are mean ±SEM, n=5-6, ***p<0.0001 vs. Control (One-way ANOVA; Tukey post-hoc test). For data plotted in graphs, see S7 Data. Scale bars: 100 µm.

**S8 Fig. Characterization of cortical phenotypes in *NexCre;CasTcKO* mice. (A-B)** Immunostaining of coronal sections for P7 control and *NexCre;CasTcKO* animals with layer makers Rorβ (**A**, red, layer IV) and Tbr1 (**B**, red, layer VI). DAPI was used to counterstain nuclei (blue). No obvert lamination phenotypes are observed. Right panels: quantification of proportion of Rorβ+ (**A**) and Tbr1+ (**B**) cells per bin. Values given are Mean ± S.E.M., n=3 animals per group, 3 sections per animal, Mann-Whitney *U* test & Bonferroni correction, no significant differences (p>0.05). For data plotted in graphs, see S6 Data. Scale bar: 500 µm.

**S9 Fig. Progenitor phenotype in E12.5 *Em×1Cre;CasTcKO cortices.* (A-B)** Detection of proliferative cells in Control and *Em×1Cre;CasTcKO* coronal sections using short-pulsed EdU at E12.5 (**A,B**, green), colabeled with the intermediate progenitor marker Tbr2 (**B,** magenta). Nuclei were counterstained with DAPI (blue). (**C**) Density of EdU+ cells/mm^2^ of cortex. p=0.34, two-tailed Mann-Whitney *U* test, n=4 independent samples per group, 3 sections per sample. (**D**) Proportion of Edu+ cells that co-express Tbr2. p=0.49, two-tailed Mann-Whitney *U* test, n=4 independent samples per group, 2-3 sections per sample. Values given are Mean ± S.E.M. For data plotted in graphs, see S11 Data. Scale bars: Lower magnification panel for A: 500 µm; Higher mag panels in A: 100 µm; B: 75 µm.

**S10 Fig. No lamination phenotypes are observed in *NexCre;β1-Integrin^loxp/loxp^* animals.** (**A-C**) Immunostaining for cortical layer markers on P7 cortical sections from Control and *Ne×Cre;β1-Integrin^lmψ/lo×ψ^* animals. Staining for Cux1 (**A**, green, layers II-III) and Ctip2 (**A**, magenta, layers V and VI), or Tbr1 (**B**, red, layer VI), or Rorβ (**C,** red, layer IV), are shown. Sections were counterstained with DAPI (blue). n=3 animals per genotype for each marker. Scale bars: 500 µm.

**S11 Fig. Some cells in mixed primary cortical cultures express RGC markers.** Example of E13.5 4DIV (four days in vitro) mixed primary cortical cultures after staining for neural markers to identify distinct cell types. Left panel: Nestin (green, neuroepithelial and radial glial cells), GFAP (magenta, astrocytes); Middle panel: Blbp (red, radial glial cells); Right panel: Tuj1 (green, postmitotic neurons), Tbr2 (magenta, intermediate progenitors). DAPI was used to counterstain nuclei (blue). n=3 independent cultures. Scale bar: 25 µm.

**S12 Fig. Cortical phenotypes on the contralateral hemispheres to the *in utero* electroporated lateral ventricles.** Immunostaining of P0 control and *Em×1Cre;CasTcKO* coronal sections of the contralateral hemispheres to the sides co­electroporated with ΔSrc + p130Cas-ZipB (negative control) or Src-ZipA + p130Cas-ZipB (FIT rescue). Sections were stained for the layer V & VI marker Ctip2 (green) and laminin (magenta), and counterstained with DAPI (blue). In utero electroporation of one side does not affect the expected phenotype on the opposite hemisphere: i.e. cobblestone phenotypes are still observed in the contralateral side of ΔSrc + p130Cas-ZipB or Src-ZipA + p130Cas-ZipB electroporated *Em×1Cre;CasTcKO* cortices. n=6 animals per genotype. Scale bar: 250 µm.

S1 Data. Numerical values for graphs in Fig 3 including Fig 3D and Fig 3E

S2 Data. Numerical values for graphs in S3 Fig including S3A Fig, S3B Fig and S3C Fig.

S3 Data. Numerical values for graph in S4 Fig.

S4 Data. Numerical values for graphs in Fig 4 including Fig 4A, Fig 4B, Fig 4C and Fig 4D.

S5 Data. Numerical values for graph in Fig 5.

S6 Data. Numerical values for graphs in S6 Fig.

S7 Data. Numerical values for graph in S7 Fig.

S8 Data. Numerical values for graphs in Fig 6 including Fig 6A, Fig 6B and Fig 6C.

S9 Data. Numerical values for graphs in S8 Fig including S8A Fig and S8B Fig.

S10 Data. Numerical values for graphs in Fig 9 including Fig 9C, Fig 9D and Fig 9E.

S11 Data. Numerical values for graphs in S9 Fig including S9C Fig and S9D Fig

S12 Data. Numerical values for graph in Fig 10.

S13 Data. Numerical values for graphs in Fig 12.

S14 Data. Numerical values for graphs in Fig 13.

## Contingency tables for cortical dysplasia penetrance

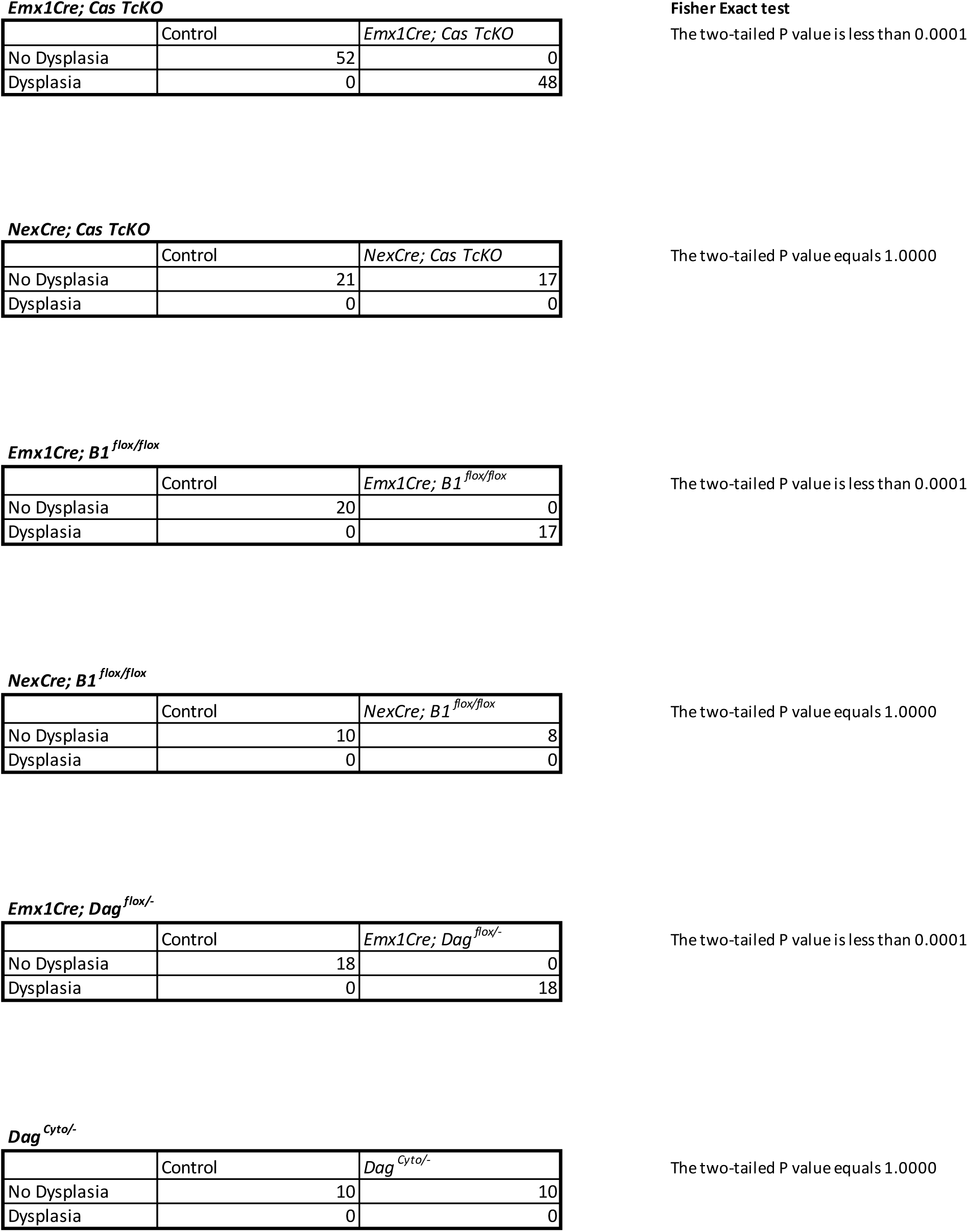

