## Supplemental figures for "An adhesion signaling axis involving Dystroglycan, β1-Integrin and Cas adaptor proteins regulates the establishment of the cortical glial scaffold"

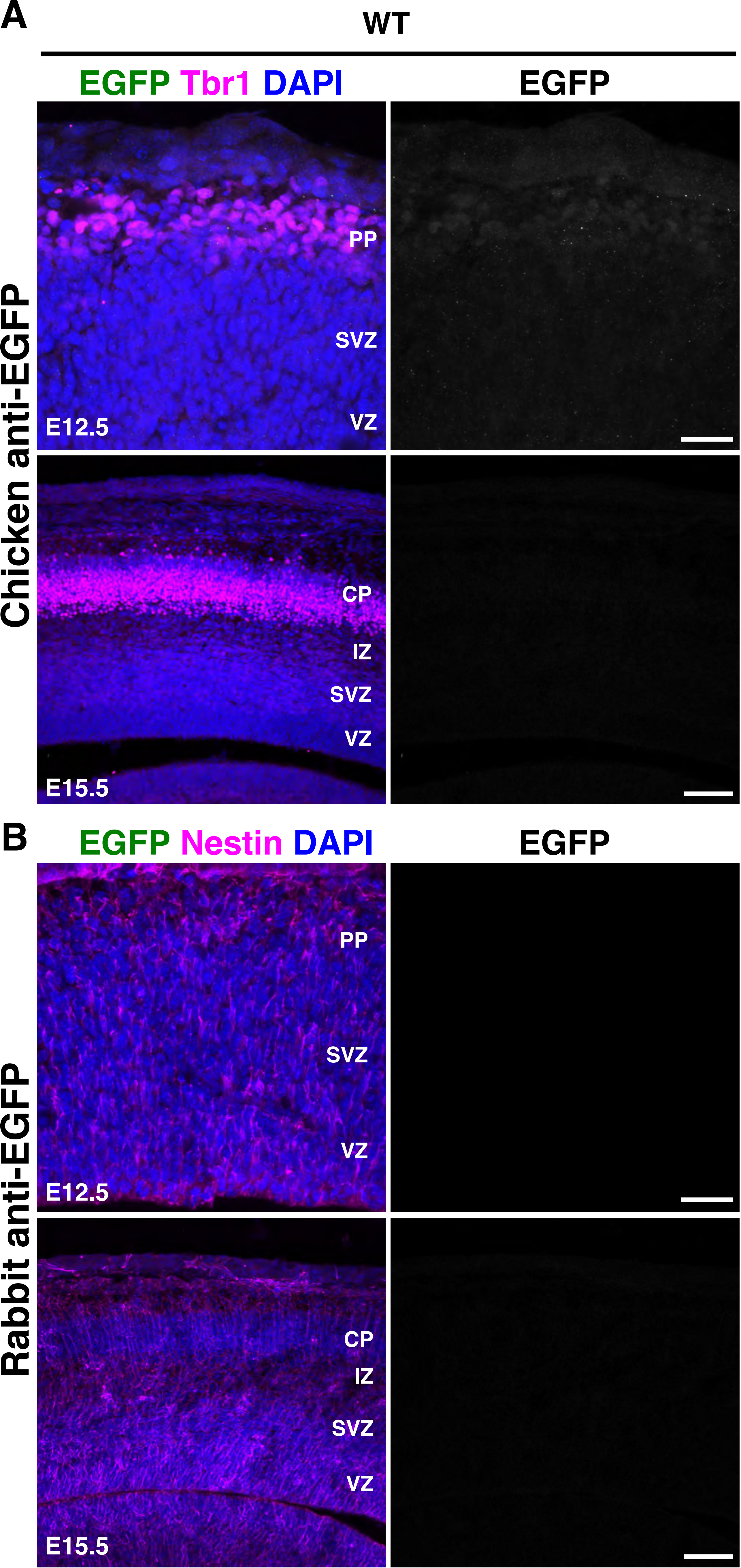

A
WT
EGFP Tbr1 DAPI
EGFP
Chicken anti-EGFP
PP
SVZ
VZ
E12.5
CP
IZ
SVZ
VZ
E15.5
B
EGFP Nestin DAPI
EGFP
Rabbit anti-EGFP
PP
SVZ
VZ
E12.5
CP
IZ
SVZ
VZ
E15.5

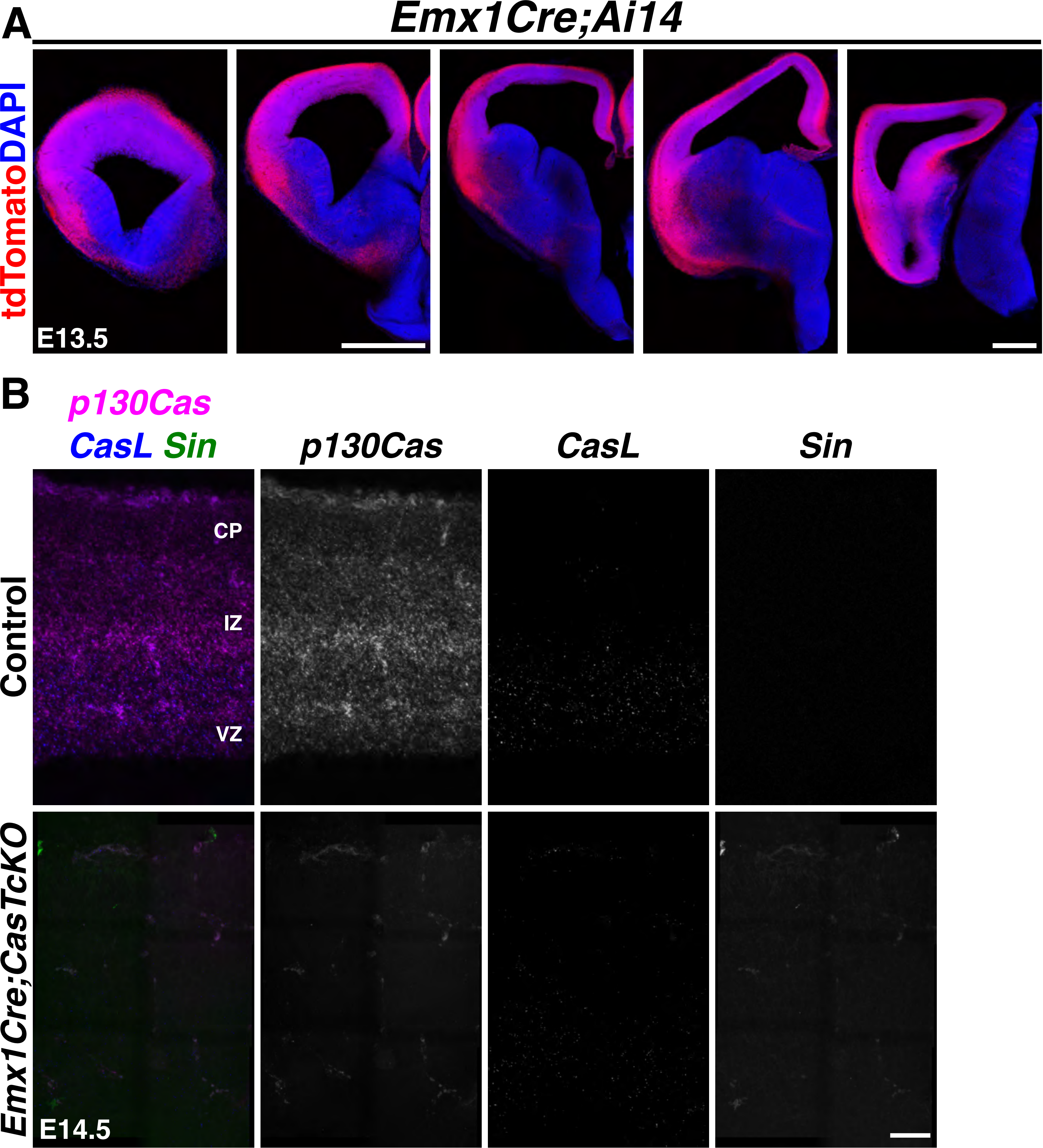
A
Emx1Cre;Ai14
tdTomatoDAPI
E13.5
B
p130Cas
CasL Sin
p130Cas
CasL
Sin
Control
CP
IZ
VZ
Emx1Cre;CasTcKO
E14.5

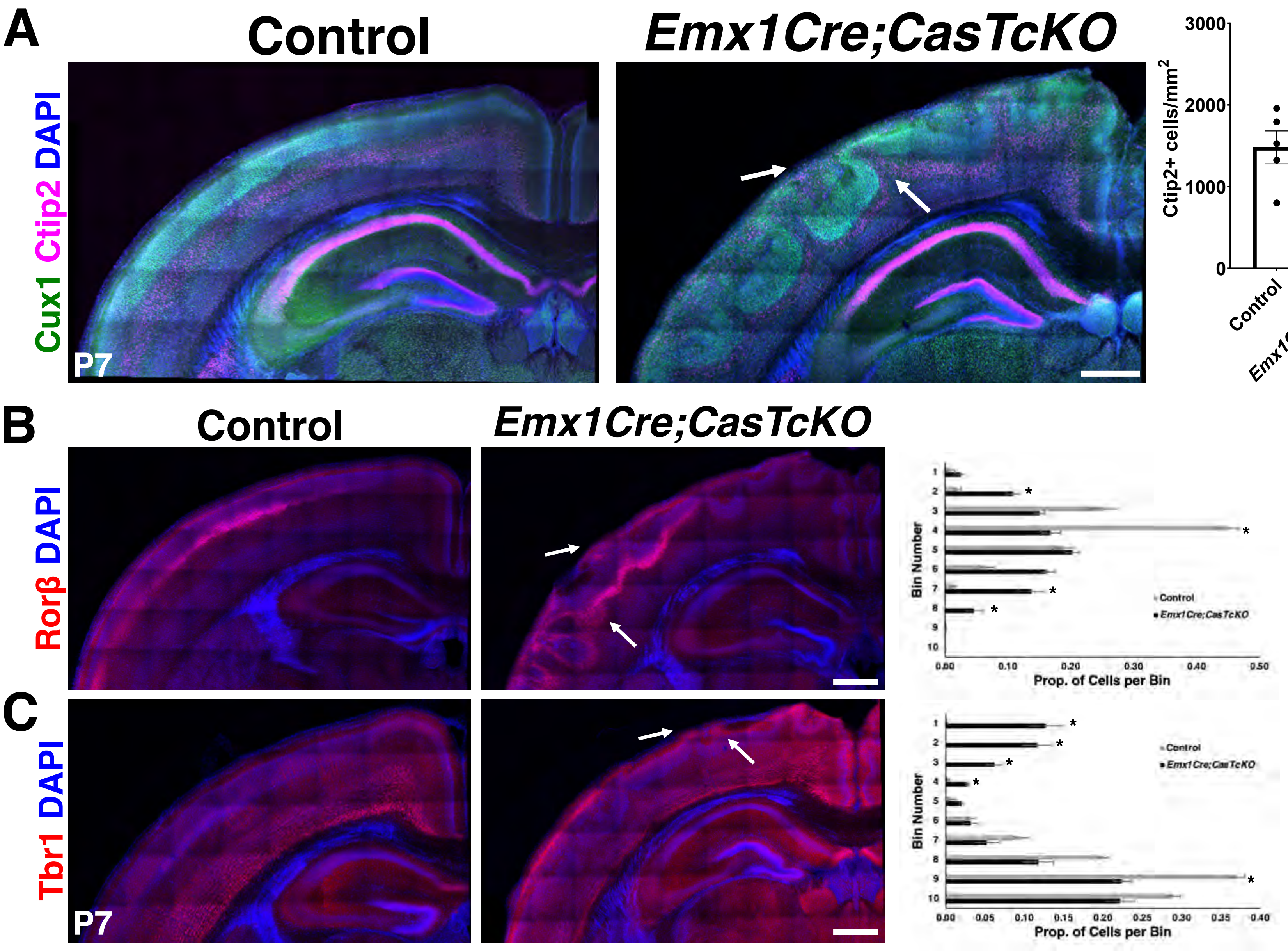
A
Control
Emx1Cre;CasTcKO
Cux1 Ctip2 DAPI
P7
Ctip2+ cells/mm2
Cux1+ cells/mm2
Control
Emx1Cre;TcKO
B
Control
Emx1Cre;CasTcKO
Rorβ DAPI
Bin Number
Prop. of Cells per Bin
Control
Emx1Cre;CasTcKO
C
Tbr1 DAPI
P7
Bin Number
Prop. of Cells per Bin
Control
Emx1Cre;CasTcKO

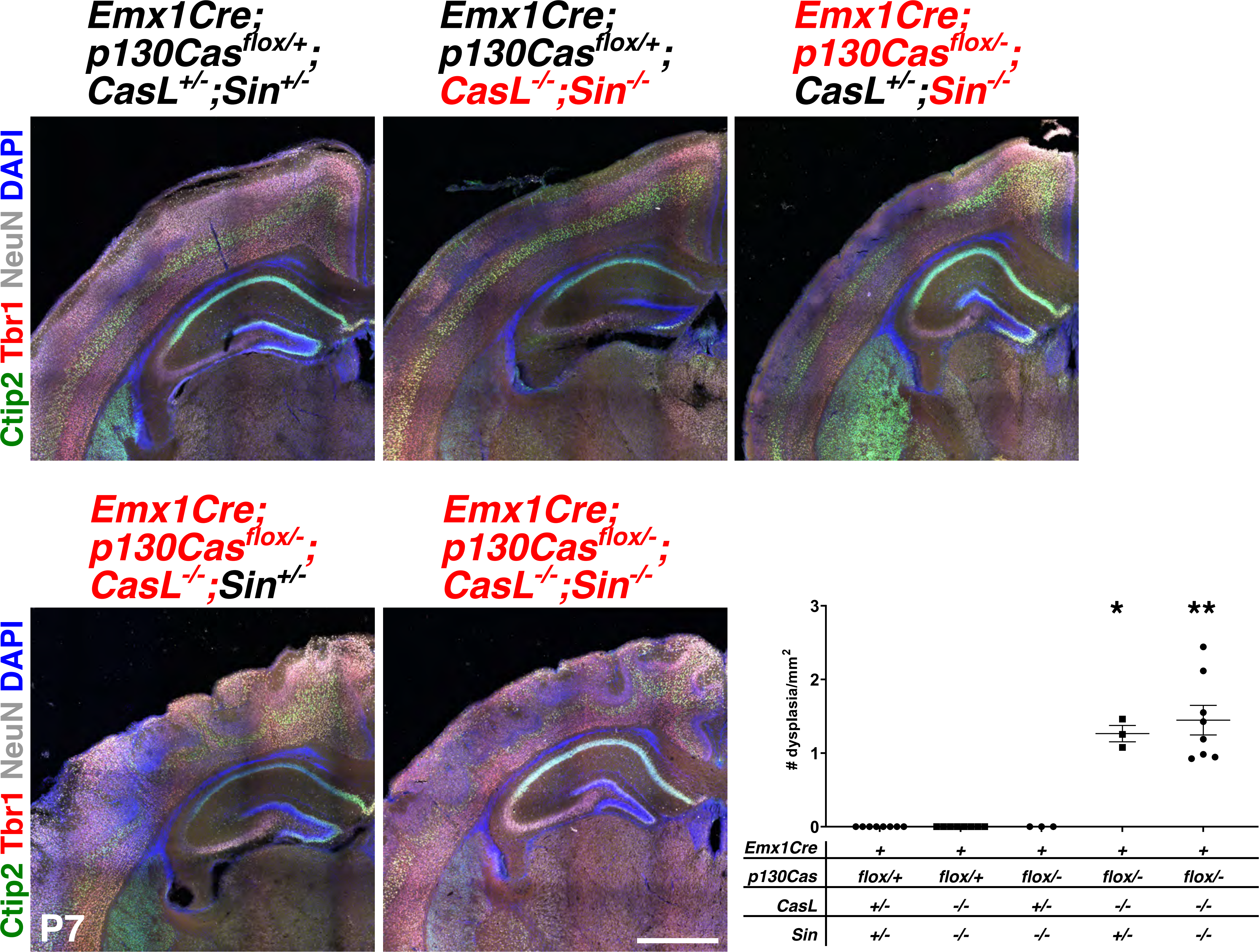
Emx1Cre; p130Cas flox/+; CasL +/-;Sin +/-
Emx1Cre; p130Cas flox/+; CasL -/-;Sin -/-
Emx1Cre; p130Cas flox/-; CasL +/-;Sin -/-
Emx1Cre; p130Cas flox/-; CasL -/-;Sin +/-
Emx1Cre; p130Cas flox/-; CasL -/-;Sin -/-
Ctip2 Tbr1 NeuN DAPI
Ctip2 Tbr1 NeuN DAPI
P7
### dysplasia/mm2
0
1
2
3
*
**
Emx1Cre + + + + +
p130Cas flox/+ flox/+ flox/- flox/- flox/-
CasL +/- -/- +/- -/- -/-
Sin +/- -/- -/- +/- -/-

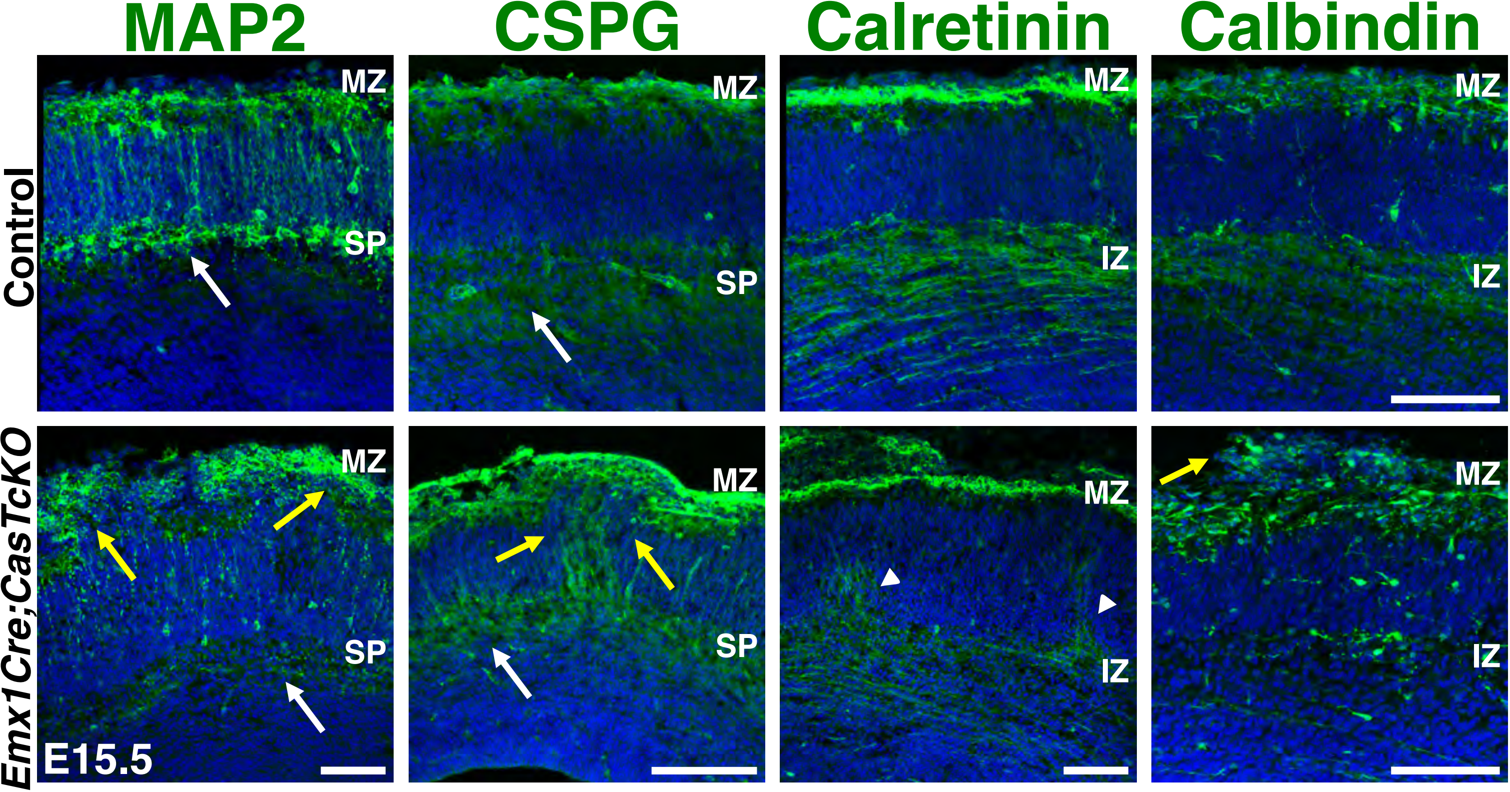
MAP2
CSPG
Calretinin
Calbindin
Control
Emx1Cre;CasTcKO
MZ
SP
IZ
E15.5

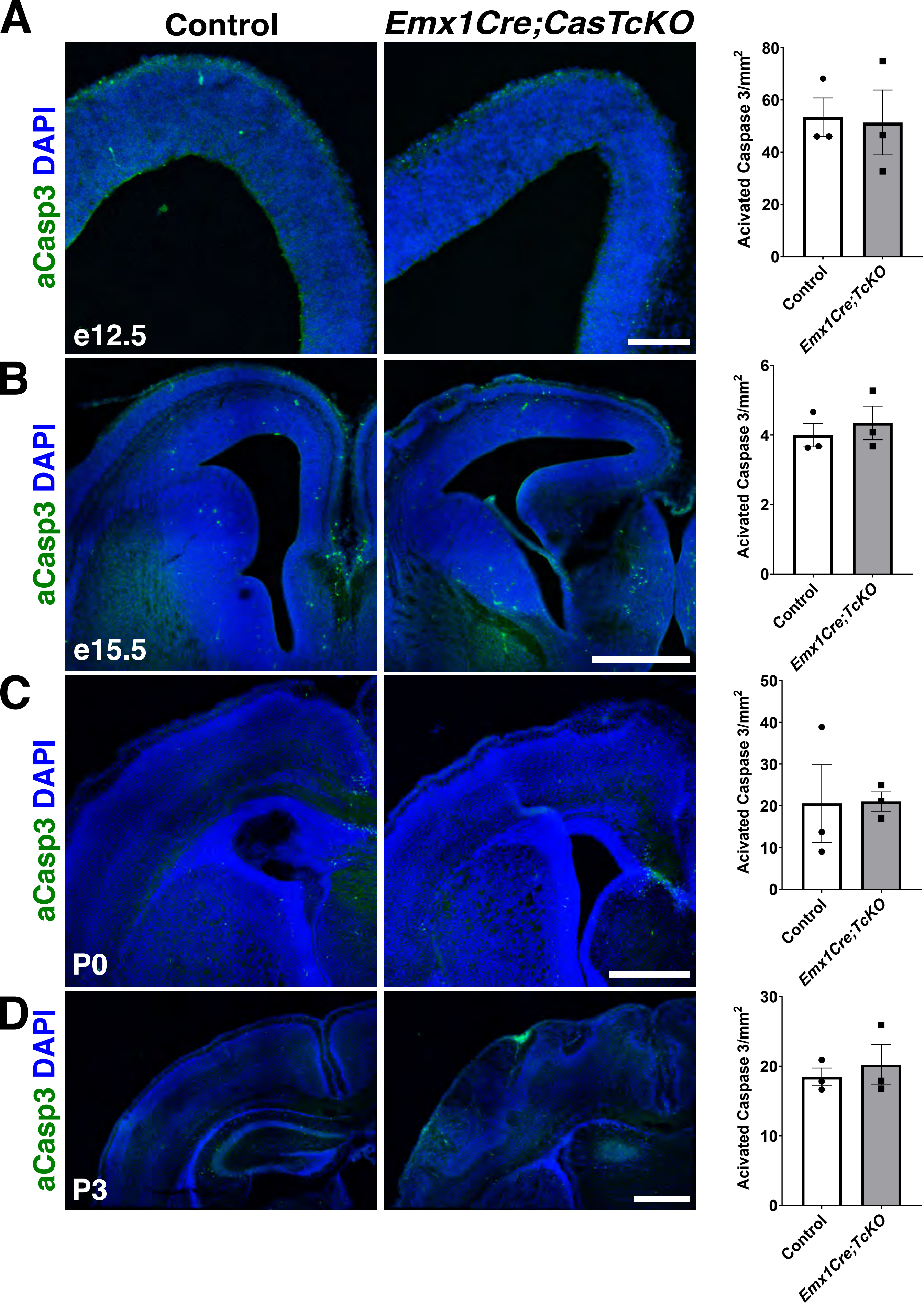
A
B
C
D
Control
Emx1Cre;CasTcKO
aCasp3 DAPI
e12.5
e15.5
P0
P3
Acivated Caspase 3/mm2
Control
Emx1Cre;TcKO
0
20
40
60
80
0
2
4
6
0
10
20
30
40
50
0
10
20
30

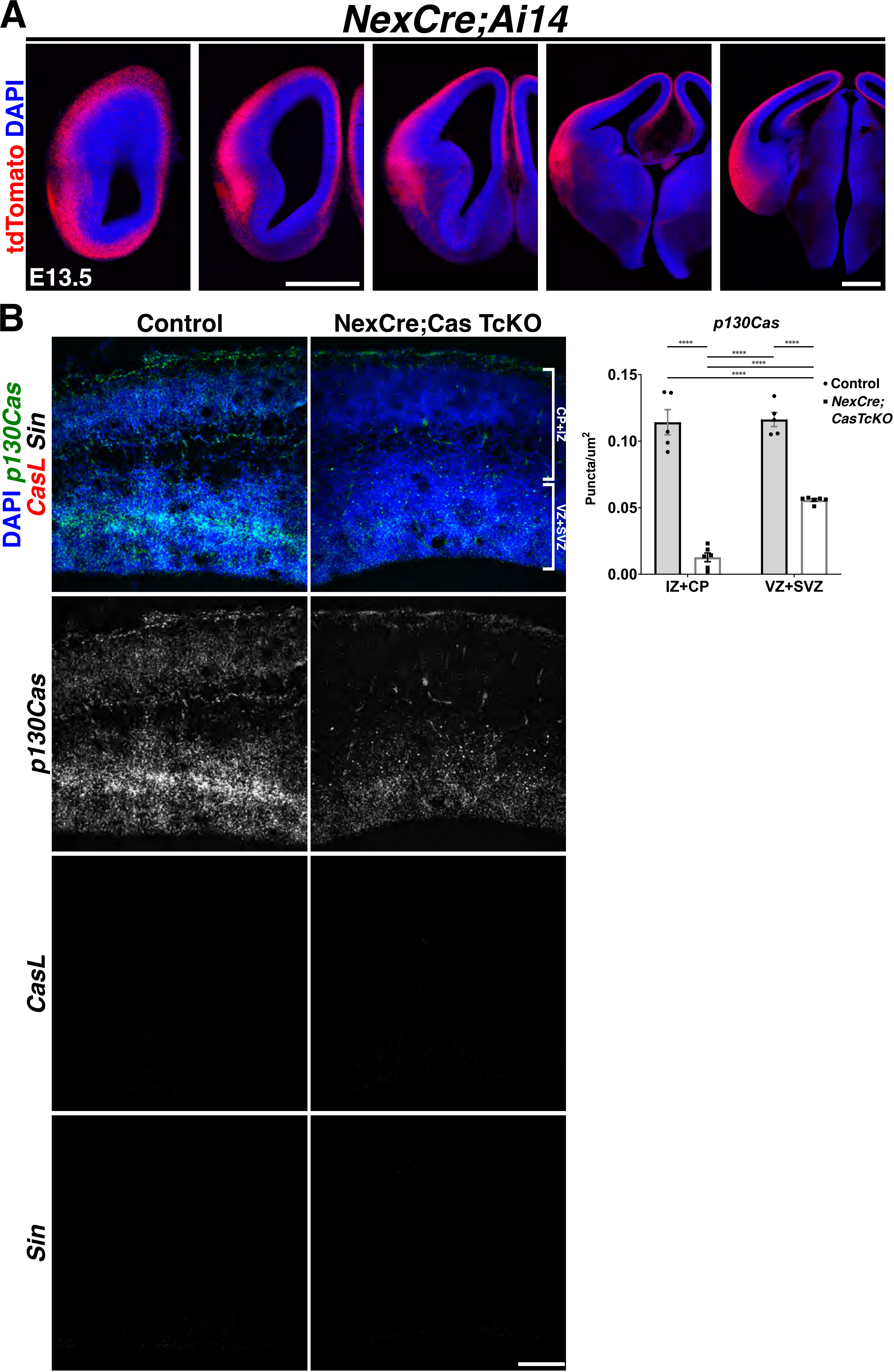

A
NexCre;Ai14
tdTomato DAPI
E13.5
B
Control
NexCre;Cas TcKO
DAPI p130Cas CasL Sin
CP+IZ
VZ+SVZ
p130Cas
CasL
Sin
p130Cas
****
0.15
0.10
0.05
0.00
Puncta/um2
IZ+CP
VZ+SVZ
• Control
▪ NexCre; CasTcKO

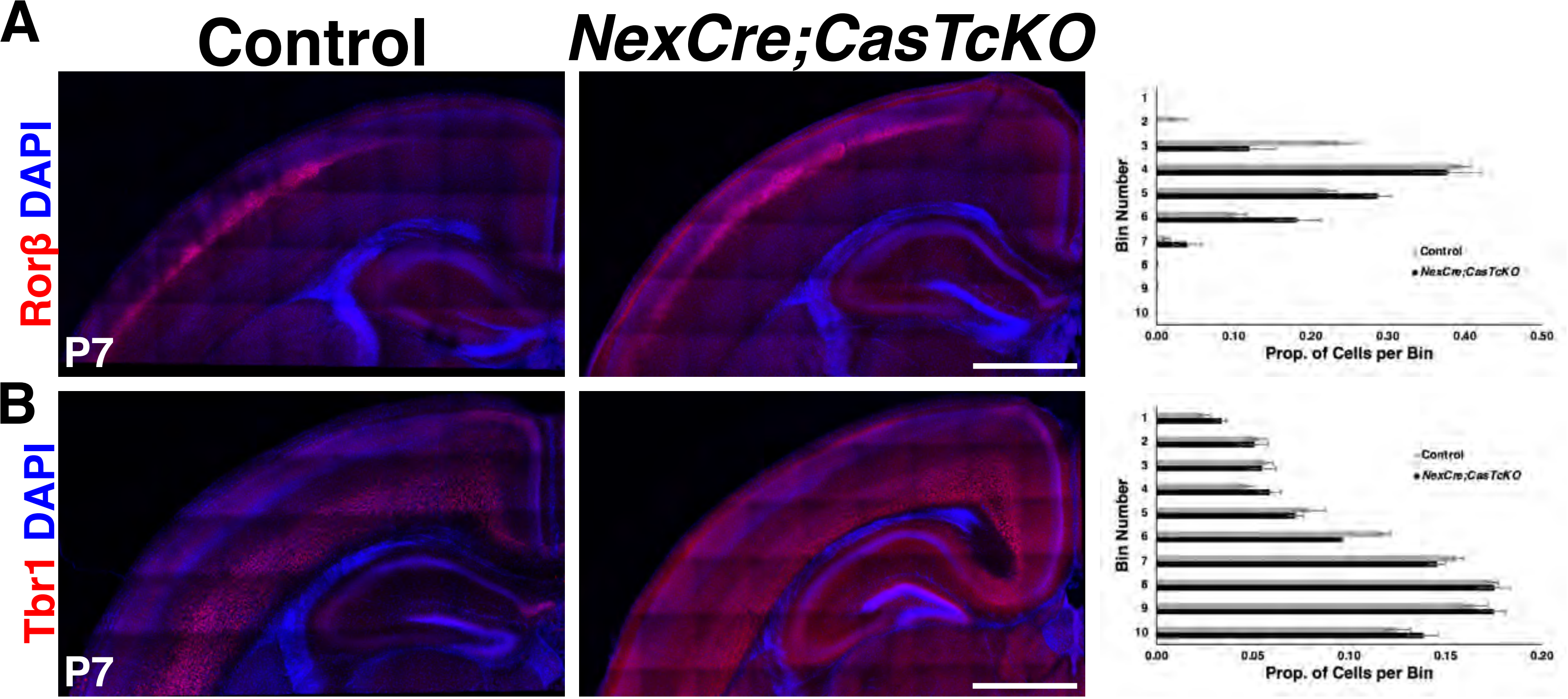

A
Control
NexCre;CasTcKO
Rorβ DAPI
P7
Bin Number
Prop. of Cells per Bin
Control
NexCre;CasTcKO
B
Tbr1 DAPI
P7
Bin Number
Prop. of Cells per Bin
Control
NexCre;CasTcKO

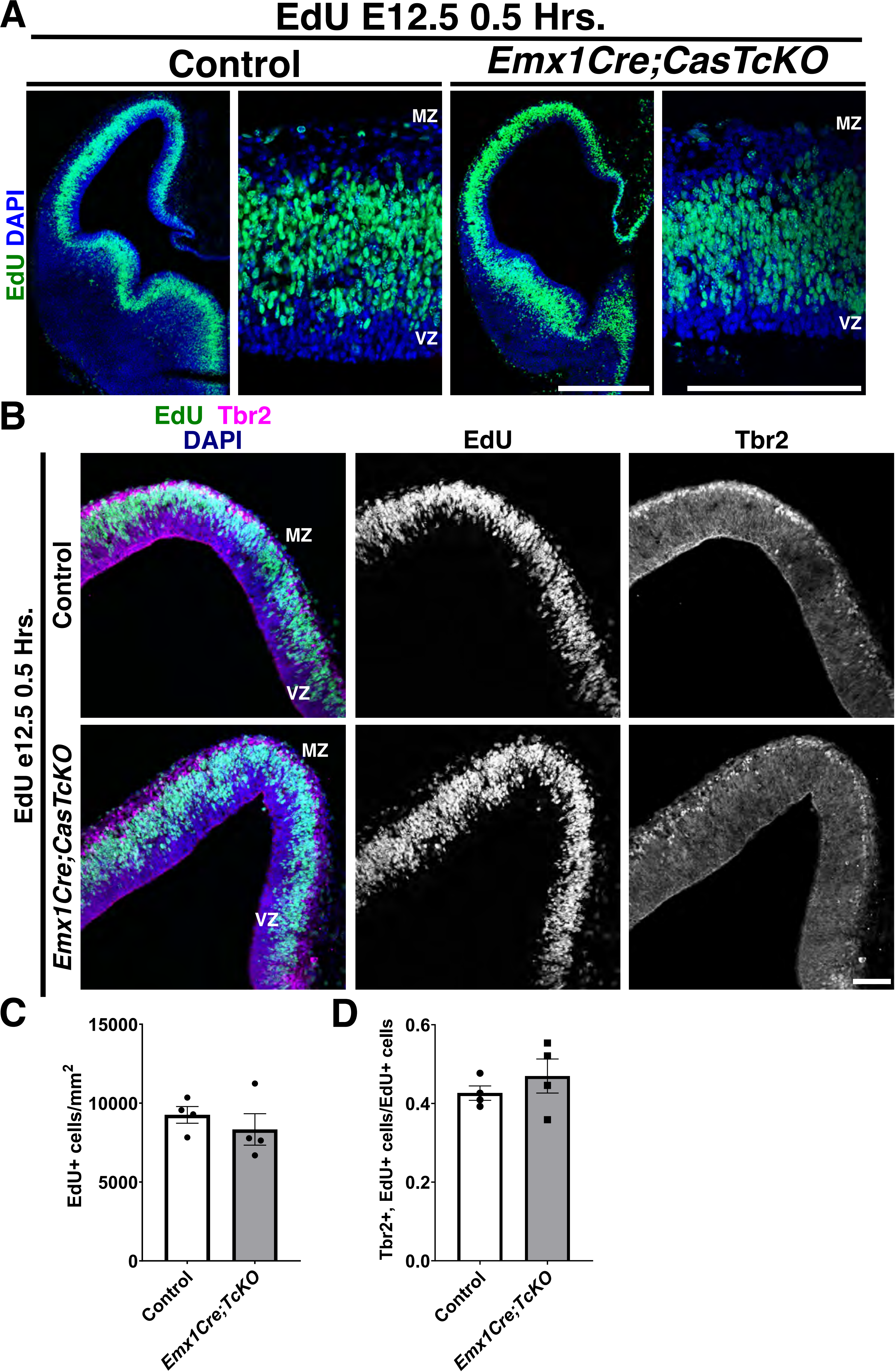

A
EdU E12.5 0.5 Hrs.
Control
Emx1Cre;CasTcKO
EdU DAPI
MZ
VZ
MZ
VZ
B
EdU Tbr2
DAPI
EdU
Tbr2
EdU e12.5 0.5 Hrs.
Control
Emx1Cre;CasTcKO
MZ
VZ
MZ
VZ
C
EdU+ cells/mm$^2$
15000
10000
5000
0
Control
Emx1Cre;TcKO
D
Tbr2+, EdU+ cells/EdU+ cells
0.6
0.4
0.2
0.0
Control
Emx1Cre;TcKO

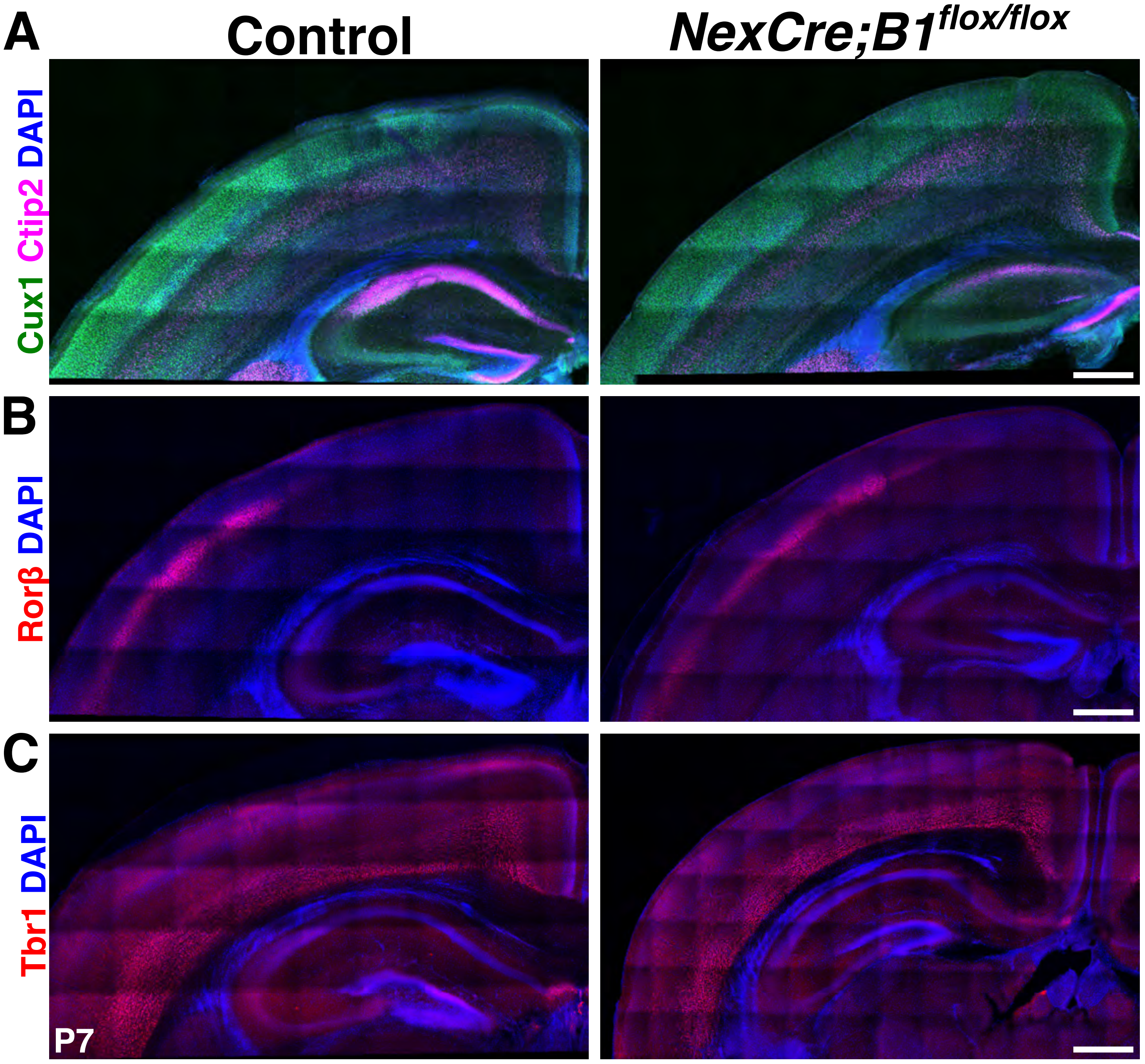

A
Control
NexCre;B1flox/flox
Cux1 Ctip2 DAPI
B
Rorβ DAPI
C
Tbr1 DAPI
P7

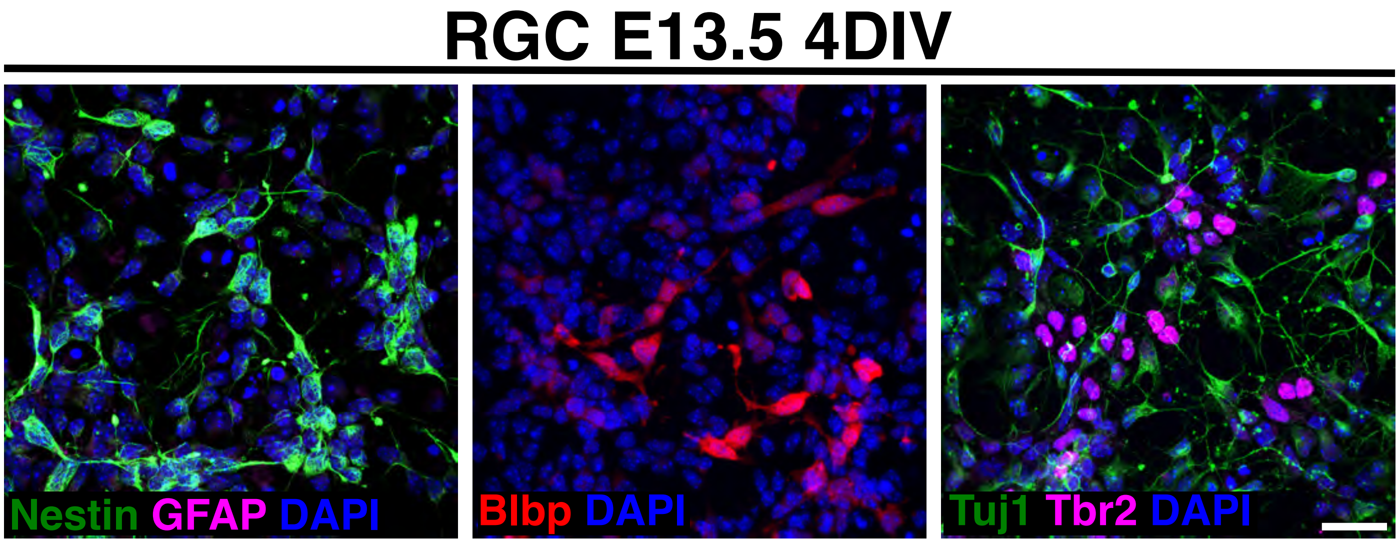
RGC E13.5 4DIV
Nestin GFAP DAPI
Blbp DAPI
Tuj1 Tbr2 DAPI

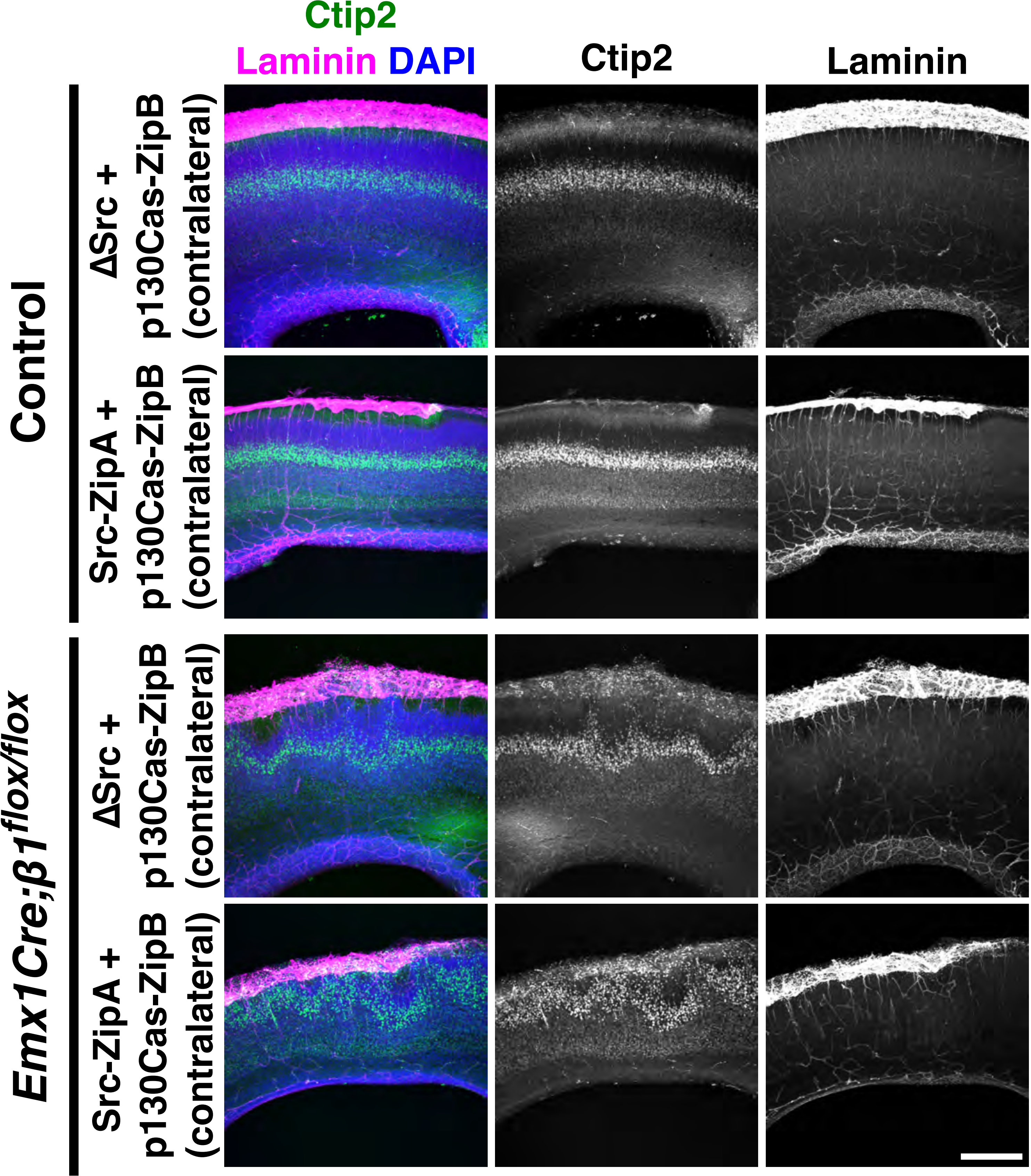

Ctip2
Laminin DAPI
Ctip2
Laminin
Control
ΔSrc +
p130Cas-ZipB
(contralateral)
Src-ZipA +
p130Cas-ZipB
(contralateral)
Emx1Cre;β1flox/flox
ΔSrc +
p130Cas-ZipB
(contralateral)
Src-ZipA +
p130Cas-ZipB
(contralateral)
